# Rescuing Ischemic Brain Injury by Rewiring Mitochondrial Electron Flow

**DOI:** 10.1101/2025.05.19.650489

**Authors:** Belem Yoval-Sánchez, Ivan Guerrero, Qiuying Chen, Sergei Sosunov, Fariha Ansari, Max Siragusa, Csaba Konrad, Zoya Niatsetskaya, Anna Stepanova, Anatoly A. Starkov, Sergei Khruschev, Jordi Magrane, Arina A. Nikitina, Oxana Bereshchenko, Ping Zhou, Liping Zhou, Marten Szibor, Ilka Wittig, Giovanni Manfredi, Steven S. Gross, Vadim Ten, Alexander Galkin

**Affiliations:** Feil Family Brain and Mind Research Institute, Weill Cornell Medicine, New York, NY, USA; Departments of Pediatrics, Robert Wood Johnson Medical School, Rutgers University, New Brunswick, USA; Department of Pharmacology, Weill Cornell Medicine, New York, USA; Department of Biophysics, Moscow State University, Moscow, Russia; Department of Brain Sciences, Faculty of Biology, Weizmann Institute of Science, Rehovot, Israel; Department of Philosophy, Social Sciences, Humanities and Education, University of Perugia, Italy; Institute for Cardiovascular Physiology, Goethe University, Frankfurt am Main, Germany; Functional Proteomics Center, Goethe University, Frankfurt am Main, Germany; Department of Cardiothoracic Surgery, Center for Sepsis Control and Care, Jena University Hospital, Jena, Germany

**Keywords:** brain ischemia-reperfusion, mitochondria, complex I, reverse electron transfer, alternative oxidase, ROS, Respiratory chain rewiring, TCA cycle

## Abstract

Mitochondrial metabolic flux alterations are critical drivers of acute ischemia-reperfusion (IR) brain injury. Reverse electron transfer (RET), defined as the upstream flow of electrons from the quinone pool to complex I, is a major source of pathological reactive oxygen species (ROS) under stress conditions. In an in vivo brain IR model, oxygen deprivation induces the buildup of RET-supporting substrates, with glycerol 3-phosphate identified as the dominant contributor in addition to succinate. Rapid oxidation of these substrates by brain mitochondria upon reoxygenation drives massive ROS production, while also leading to over-reduction and dissociation of the complex I flavin mononucleotide (FMN) cofactor. The resulting FMN-deficient complex I becomes catalytically impaired, unable to oxidize NADH or to produce ROS.

To mitigate RET and preserve complex I function, we used transgenic mice xenotopically expressing alternative oxidase (AOX). This enzyme bypasses complexes III and IV by directly oxidizing the reduced quinone pool and passing electrons onto molecular oxygen. AOX expression did not alter complex I abundance, supercomplexes assembly, or basal respiration rates, but effectively diverted electrons from the quinone pool, decreasing RET flux via complex I and limiting ROS generation during IR. For the first time we showed that AOX expression and attenuation of RET preserved complex I FMN binding, suppressed oxidative stress, and conferred neuroprotection *in vivo*. Our findings reveal a novel strategy for rewiring mitochondrial electron flux to mitigate initial IR brain injury, highlighting modulation of the quinone pool by AOX as a potential therapeutic strategy for IR.

## Introduction

Ischemia-reperfusion (IR) brain injury is a leading cause of mortality and long-term neurological disability in adults following ischemic stroke and neonates with hypoxic-ischemic encephalopathy. The economic burden is substantial, with lifetime care costs reaching millions of dollars per patient in the U.S.^1,2^. Given the brain’s reliance on mitochondrial oxidative phosphorylation for ATP production, redox and ion balance, and signal transduction, mitochondrial dysfunction emerged as a key driver of IR pathology.

In the process of oxidative phosphorylation, electrons from flavin-dependent dehydrogenases – complex I (NADH:ubiquinone oxidoreductase), complex II (succinate dehydrogenase), and glycerol 3-phosphate dehydrogenase (GPDH) – are transferred to the ubiquinone pool and subsequently passed through complexes III and IV to molecular oxygen^3^. The coupling of electron transfer to proton translocation establishes the proton-motive force that drives ATP synthesis. However, during ischemia, oxygen deprivation halts electron flow at complex IV, leading to respiratory chain reduction, ATP depletion, and metabolic substrates accumulation. In both mice and humans, oxygen levels decline within seconds of ischemia, with ATP depletion following shortly after^4-6^. Reperfusion restores mitochondrial respiration but also can trigger oxidative stress and enzyme damage.

Mitochondrial complex I is particularly vulnerable to IR injury^3,7^ and a major contributor to mitochondrial reactive oxygen species (ROS) production^8,9^. Multiple *in vivo* studies have reported a decline in complex I activity in the brain post-IR^10-19^, yet the underlying mechanisms remain incompletely understood. We and others have recently demonstrated that IR results in flavin mononucleotide (FMN) dissociation from complex I, impairing its function^17-19^, but the mechanisms of this process have not been elucidated.

Complex I, a multi-subunit enzyme, is central to mitochondrial metabolism, catalyzing electron transfer from NADH to ubiquinone, therefore regenerating NAD^+^ for the TCA cycle and glycolysis^3,20^. Complex I is composed of 45 protein subunits, along with a non-covalently bound flavin mononucleotide (FMN), and eight iron-sulfur (Fe-S) clusters. Notably, under conditions of a highly reduced quinone pool and sufficient proton-motive force, complex I can operate in reverse, driving electrons upstream to reduce NAD^+^ in a process known as reverse electron transfer (RET). It should be stressed that, the entire pool of matrix pyridine nucleotides can be rapidly reduced by RET *in situ*. Therefore, during the steady-state, RET via complex I may be redirected to an alternative acceptor, such as molecular oxygen driving massive production of ROS upon reperfusion. RET was first described in mitochondria that were oxidizing succinate or glycerol 3-phosphate^21,22^, and has recently been implicated in pathological ROS generation during IR^23-27^.

In the present report, we identified a metabolic shift favoring RET substrate accumulation during ischemia, leading to excessive complex I reduction and FMN dissociation upon reperfusion. Using transgenic mice expressing alternative oxidase (AOX), which bypasses complexes III and IV by directly oxidizing the quinone pool, we demonstrate attenuation of RET, prevention of functional loss of complex I, and neuroprotection following IR. These findings establish RET as the main driver for FMN dissociation from complex I and the fundamental pathological mechanism that underlies stroke and hypoxic-ischemic brain injury, positioning AOX as a potential therapeutic strategy.

## Results

### Effect of brain IR on mitochondrial respiratory chain enzymes

In this study, we employed the Rice-Vannucci model of regional hypoxia-ischemia (HI) brain injury in neonatal mice (Fig. 1a)^28,29^. HI insult led to a substantial infarction of the ipsilateral hemisphere, with an average infarct volume of 44.4 ± 20.5%, as assessed by TTC staining (Fig. 1b, c). Given the dependence of both oxidative phosphorylation and mitochondrial ROS production on oxygen availability^8^, we performed real-time measurements of brain oxygenation *in vivo* by the phosphorescence quenching method using Oxyphor PdG4 as a probe^30^. Real-time data presented in Fig. 1d show measurements of brain oxygen at the hemisphere ipsilateral to the carotid artery ligation side during ligation, hypoxic exposure, and reoxygenation. Notably, changes in inhaled oxygen were closely mirrored by fluctuations in brain oxygen levels (Fig. 1e). Systemic circulatory arrest resulted in complete anoxia (Fig. 1e), while in our model of HI we detected a presence of residual oxygen of 4-5 µM in the ipsilateral hemisphere.

**Figure 1.**
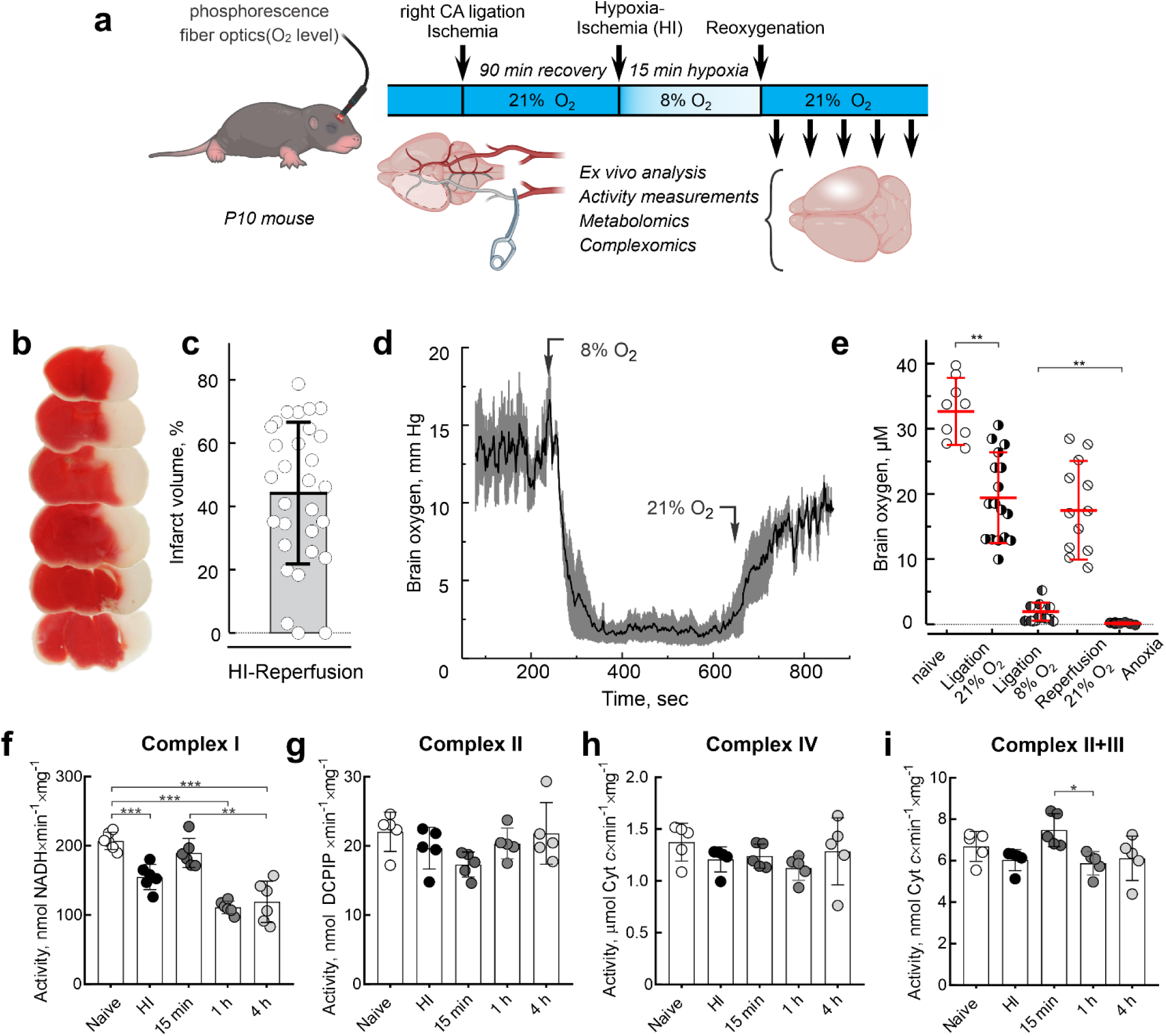
Mitochondrial complex I is impaired after brain IR. **a,** Diagram of neonatal HI-reperfusion model of *in vivo* brain injury. **b,** TTC-staining of coronal brain sections 24 h after HI-reperfusion. Functional tissue was stained (red), and the infarcted tissue was not stained (white). **c,** Infarct volume is shown as a percentage of the hemisphere volume. **d,** A representative trace showing oxygen level change in the expected infarct area in the right hemisphere of P10 mice with occluded right carotid artery during hypoxic treatment with following reoxygenation (8 and 21% oxygen, respectively) during *in vivo* HI-reperfusion experiment. Grey error bars represent SEM based on 3-4 biological replicates. **e,** Oxygen level in the affected area in the right hemisphere in naïve animals (−), mice with right carotid artery ligated (4), during HI at 8% oxygen (6) and reperfusion (<S), or after euthanasia (complete anoxia, ,). Data are means ± s.d., n=12-17. **f-i,** enzymatic activities of respiratory complexes measured in mitochondrial membranes prepared from ipsilateral hemisphere immediately after HI and at different times after reperfusion. **f,** NADH:ubiquinone reductase of complex I; **g,** succinate:DCIP reductase activity of complex II; **h,** ferrocytochrome *c* oxidase activity of complex IV; **i,** succinate:ferricytochrome *c* combined activity of complexes II and III. Mitochondrial membranes were isolated from the brain samples after HI and activity measured as described in the Materials and Methods section. Data are means ± s.d., n=5–6 per group. * p<0.05, ** p <0.01, *** p<0.001, one-way ANOVA with Dunnett’s test.

Next, we assessed the functional activity of individual respiratory chain complexes in mitochondria isolated from the ipsilateral hemisphere immediately after HI and at various time points post-reperfusion. Consistent with previous findings^10-19^, brain IR resulted in a significant decrease in mitochondrial complex I activity, whereas the activities of complexes II, IV, and II+III remained largely unchanged within the first 4 h of reperfusion (Fig. 1 f-i).

### Brain metabolomics profiling after IR

Mitochondria rapidly respond to fluctuations in oxygen availability, modulating key catabolic pathways. To characterize the metabolic signature of brain oxygen deprivation, we performed high-resolution liquid chromatography-tandem mass spectrometry (LC-MS) of brain tissue from the ipsilateral hemisphere following *in vivo* IR. A global overview of metabolite changes in HI relative to normoxia presented as a volcano plot highlights numerous differently abundant metabolites (Fig. 2a). Uniform manifold approximation and projection (UMAP) analysis demonstrates a clear pattern of metabolite changes during HI and at different time points of reperfusion (Fig. 2b). ATP and ADP levels (Fig. 2c, d), along with other nucleotide triphosphates and diphosphates (*e.g.*, GTP/GDP, not shown), declined sharply within 15 min of IR, coinciding with phosphocreatine depletion (Fig. 2j). Concurrently, adenylates and purine nucleotides underwent progressive degradation to nucleosides, nucleobases, and ultimately xanthine (Fig. 2g-i) as shown before^26,31-33^. Notably, ATP and total phosphoadenylates (ATP+ADP+AMP, Fig. 2f) failed to recover to normoxic levels even 4 h post-reperfusion, indicating a sustained energy deficit and significant depletion of adenylate pool. This suggests that, during early reperfusion, an insufficient ADP supply limits ATP resynthesis by oxidative phosphorylation. In parallel, we observed a pronounced accumulation of respiratory substrates, including succinate and glycerol 3-phosphate, which persisted for at least 15-30 min post-reperfusion before returning to baseline (Fig. 2k, l). Postischemic elevation and oxidation of these substrates has been associated with RET and excessive ROS production, with complex I serving as a major site of ROS generation^8,16,17,23,34-39^. Accordingly, we detected elevated levels of cysteic acid (Fig. 2m), a known marker of oxidative protein damage^40,41^. We recently demonstrated that RET induces over-reduction of the FMN redox center in complex I, leading to dissociation of the flavin cofactor^17-19^. While our metabolic analysis did not reveal significant changes in tissue FMN levels (not shown), we detected a striking increase in free riboflavin concentration (Fig. 2n). This likely reflects FMN hydrolysis by intramitochondrial FMN phosphatase^42,43^, leading to elevated riboflavin levels shortly after HI, that return to the basal level 4 h after reperfusion.

**Figure 2.**
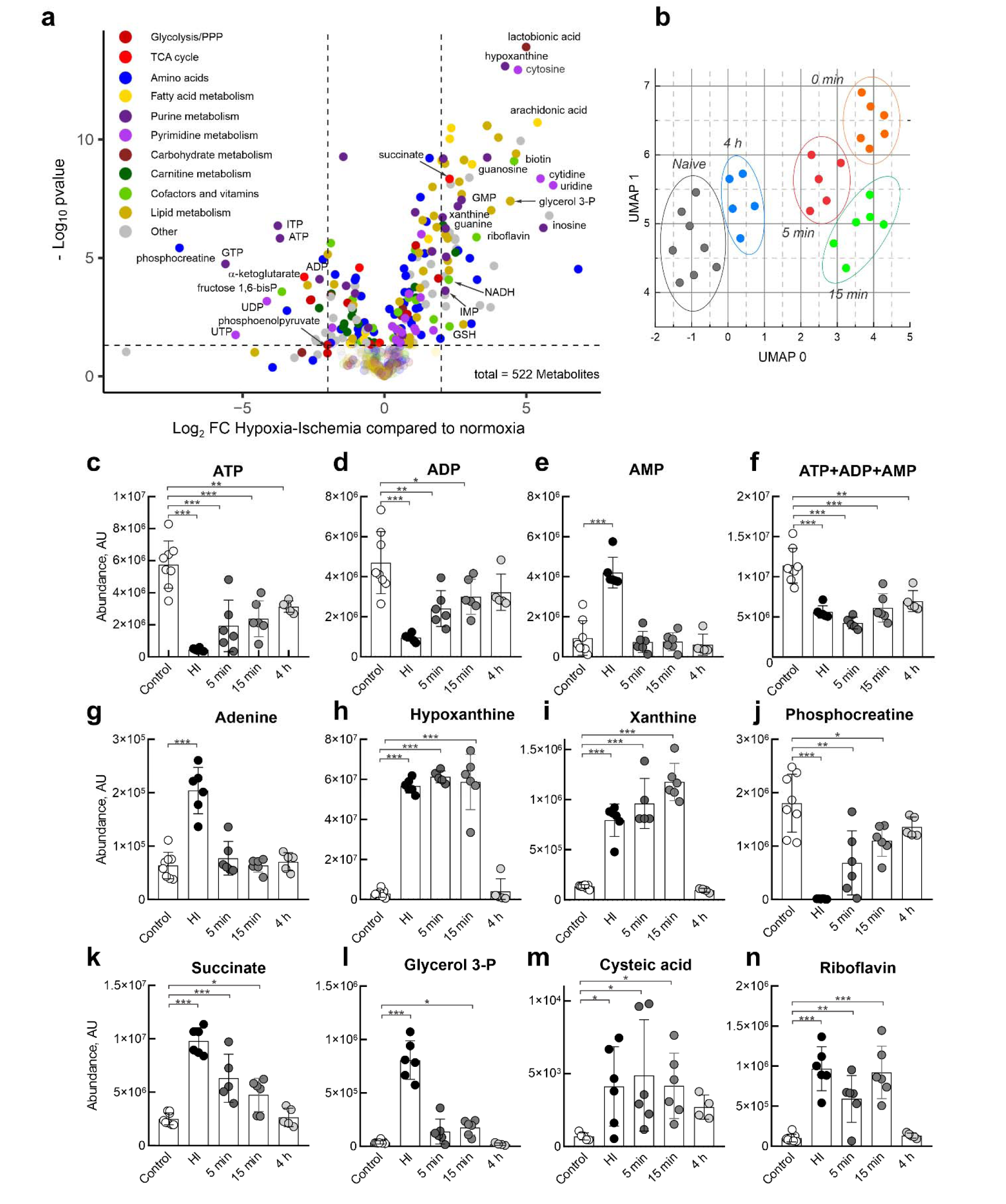
Metabolic changes in the brain following IR. **a,** Volcano plot showing fold changes (log_₂_ FC) in metabolite abundance in the ipsilateral hemisphere after HI compared to normoxia, with statistical significance (-log_₁₀_ p-value) from untargeted LC-MS analysis. Each dot shows a metabolite, color-coded by metabolic pathway, and represents the mean of 6-8 replicates. **b,** Uniform manifold approximation and projection (UMAP) of metabolic profiles, showing distinct clustering of samples based on time points: naïve (black), 0 min (orange), 5 min (red), 15 min (green), and 4 h (blue) post-reperfusion. Metabolites are listed in Supplementary Table 1. **c-n,** Temporal dynamics of selected metabolites following IR. Levels of **c**, ATP; **d**, ADP; **e**, AMP; **f**, total phosphoadenylates (ATP+ADP+AMP) show a rapid decline post-HI, with incomplete recovery at 4 h. **g–i,** Breakdown of adenylates leads to increased adenine, hypoxanthine, and xanthine. **j,** Phosphocreatine depletion reflects impaired energy homeostasis. **k–l,** Accumulation of succinate and glycerol 3-phosphate suggests altered mitochondrial respiration. **m,** Increased cysteic acid, a marker of oxidative protein damage. **n,** elevated riboflavin levels indicate flavin dephosphorylation. Data are means ± s.d., n=5–8 per group, *p<0.05***, **p <0.01, ***p<0.001, one-way ANOVA with Dunnett’s test.

### MALDI Imaging

To analyze metabolites in the ipsilateral hemisphere using LC-MS/MS, we prepared extracts from the tissue excised from the respective infarct location for comparison with extracts obtained from sham control animals. Notably, there is always a risk that non-affected tissue constitutes a significant fraction of the material obtained from the assumed site of the infarct. In addition, experimental mice are subjected not only to artery occlusion, but also to hypoxic exposure, therefore, it is most informative to compare similar areas of ipsi- and contralateral hemispheres or specific brain subregions from the same animal. To obtain a highly accurate distribution and relative quantification of various metabolites across both brain hemispheres, we employed MALDI Imaging analysis of the frozen brain slices to assess the abundance of metabolites that contribute to glycolysis, the TCA cycle, fatty acid oxidation, pathways of nucleotide and amino acid transformation, along with the distribution of vitamins and inorganic species (Fig. 3). The ischemic region was clearly identified based on the heat maps of expected metabolite abundances (Fig. 3a). By analyzing the metabolite distribution and employing image segmentation techniques, partial least-squares discriminant (PLS-D) analysis, and unsupervised bisecting k-means clustering, we were able to confidently discriminate the injured and healthy regions in the mouse brain subjected to HI. PLS-D analysis showed separation of pixels in ipsi- vs. contra-lateral hemispheres indicating a significant difference in the metabolites in the affected compared tissue to the non-affected tissue (Extended Data Fig. 2). Using a home-made plugin to the *napari* software platform (Supplementary Data 2), we could define two symmetrical regions of interest in the left and right brain hemispheres and plot volumes of distribution for each metabolite as split violin plots comparing these regions (Fig. 3 b-d). Notably, we found several key metabolites that were significantly different after HI in the ipsi- and contralateral hemispheres, clearly delineating the location of a future infarct (Fig. 3 and Extended Data Figure 2).

**Fig. 3.**
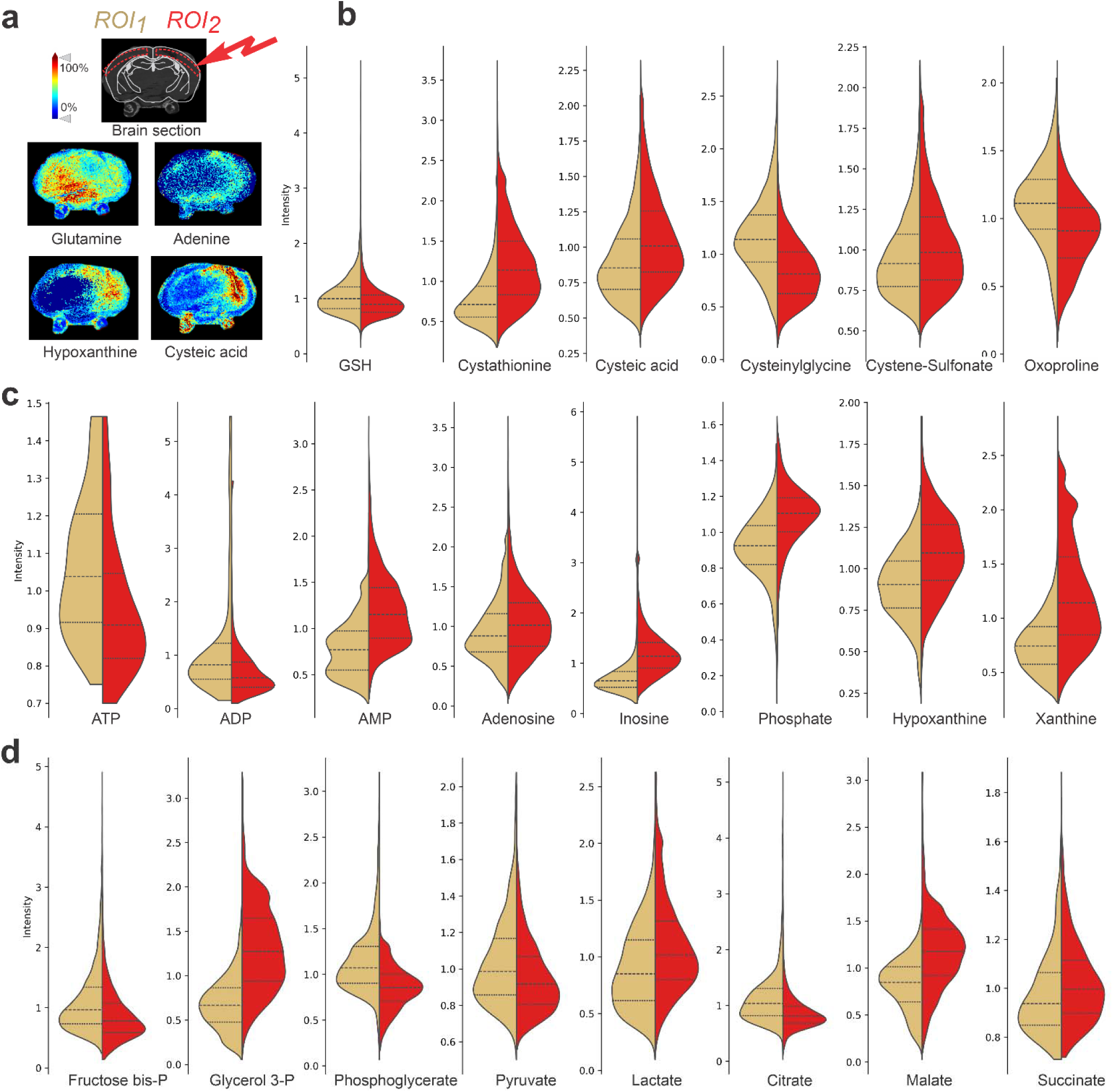
Spatial distribution of metabolites in the brain after IR by MALDI Imaging. **a,** Representative images of individual metabolites distribution in the brain slices after HI obtained by *in situ* MALDI imaging. Mouse heads after HI were immediately snap-frozen. Sectioned brain slices at 10 μm thickness were pretreated and then used for *in situ* imaging. Mass imaging data were acquired in negative ionization mode with 80 μm spatial resolution. **b-d,** Split violin plots showing the pixel intensity distribution of individual metabolites in the left and right hemispheres (ROI1 yellow and ROI2 red, respectively). **b,** Cysteine oxidation. **c,** Adenylate nucleotides degradation. **d,** Glycolysis and TCA cycle. Combined pixel distribution using the values from 2-3 brain sections from three different biological replicates was used. In all graphs corrected p-value was less than 0.001, unpaired two-tailed t-test.

Overall, MALDI imaging metabolomics data of the cortical tissue after oxygen deprivation confirm results from brain extracts analyses (Fig. 2), showing accumulation of lactate and ischemia-associated increases in succinate and glycerol 3-phosphate, as well as degradation of adenylates and other purine nucleotides in the ipsilateral right hemisphere. Remarkably, observed low levels of citrate, which provides an antioxidant action by chelating metal ions, can make this region more vulnerable to ROS damage (Fig. 3d). Increased level of oxidative stress was further evident from a dramatic increase in cysteic acid and cystathionine in the injured area of the right hemisphere (Fig. 3b). In addition, glutamine synthesis from glutamate may be impaired due to energy failure, as glutamine synthase requires ATP for activity. Glutamate levels were also decreased consistent with a low oxoproline level (Fig. 3b), potentially explaining low levels of glutathione and its breakdown product cysteinylglycine. (Fig. 3b).

### RET induces FMN dissociation from mitochondrial complex I

RET-supporting substrates, succinate, and glycerol 3-phosphate, accumulate during oxygen deprivation in brain tissue, and are subsequently oxidized within the first 30 min of reperfusion (Fig. 2 and 3). To examine the effect of these substrates on mitochondria *in vitro*, we incubated isolated intact brain mitochondria for 20 min in conditions of either reverse (succinate, glycerol 3-phosphate or both substrates together) or forward (malate and pyruvate) electron transfer (Fig. 4). Unlike oxidation of malate/pyruvate, RET substrates supported a much higher rate of H_2_O_2_ production (Fig. 4a-d). RET resulted in a gradual decay in the rates of ROS generation (Fig. 4a-d), with the strongest effect observed when succinate and glycerol 3-phosphate were combined (Fig. 4e-f). No significant change in the H_2_O_2_ production rate was observed during malate and pyruvate oxidation (Fig.4f, grey bars). We next investigated whether the mechanism of complex I inactivation and RET-dependent ROS generation decline is linked to the dissociation of complex I natural cofactor FMN. After separation of membrane complexes by high resolution Clear Native (hrCN) PAGE, we quantified complex I FMN by fluorescent scanning^44^ (Fig. 4h). We found that the shortest half-time of H_2_O_2_ release rate decay (Fig. 4e), the highest degree of complex I inactivation (Fig. 4g), and the lowest residual complex I FMN content (Fig. 4i) were observed when mitochondria oxidize a combination of two RET substrates: succinate and glycerol 3-phosphate. Another mitochondrial enzyme potentially able to undergo reduction-induced dissociation of non-covalently bound FMN is dihydroorotate dehydrogenase. Given the absence of any detectable effect of RET on dihydroorotate dehydrogenase activity (Extended Data Fig. 3), complex I-bound flavin emerges as the predominant target of RET.

**Figure 4.**
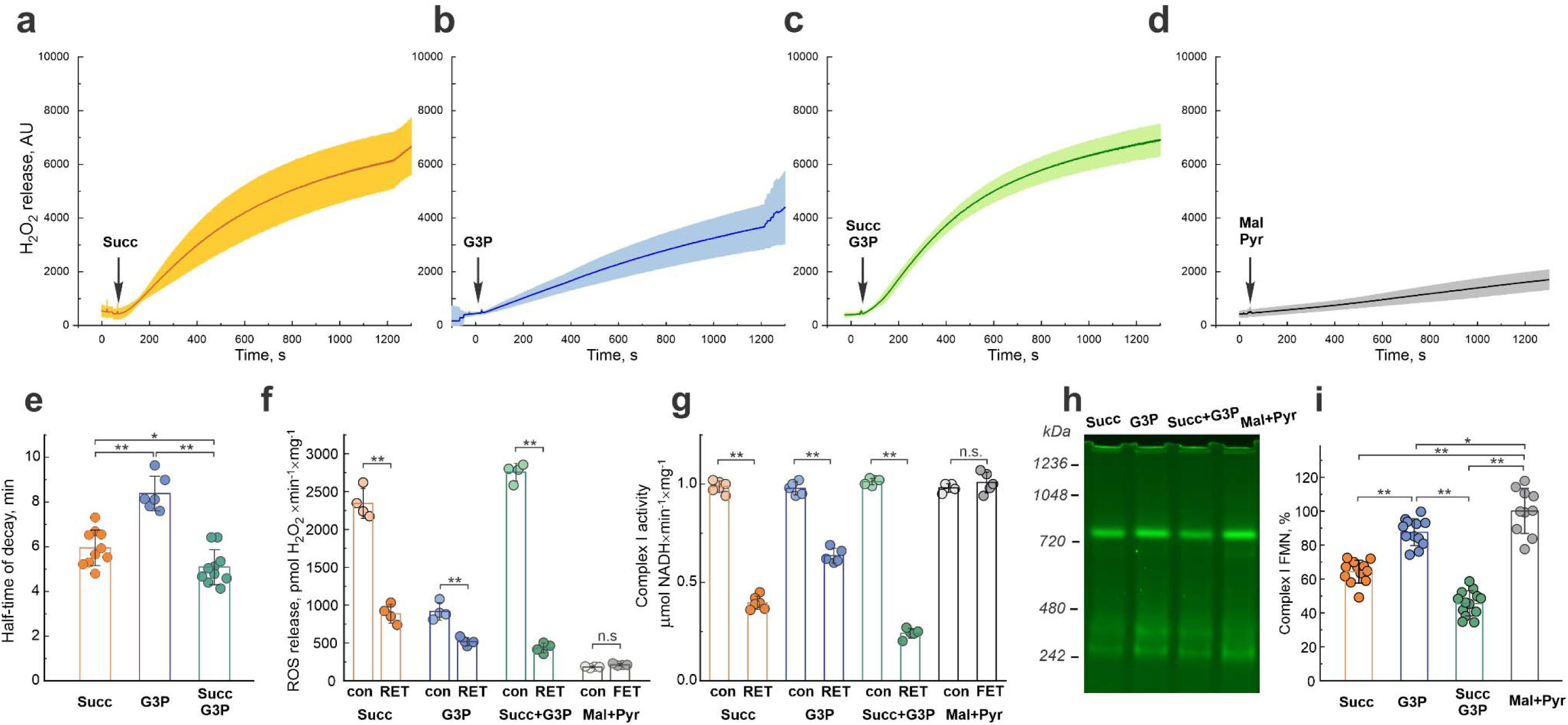
Effect of RET on mitochondria. **a-d**, Decline of ROS formation during oxidation of various substrates by intact brain mitochondria: **a,** succinate; **b,** glycerol 3-phosphate; **c,** succinate and glycerol 3-phosphate; **d,** malate and pyruvate. Addition of substrate is shown by arrow. Error bars represent one standard deviation based on triplicate measurements of different mitochondria isolations. **e,** Comparison of the half-time of the H_2_O_2_ release rate decay in mitochondria oxidizing different substrates. **f,** H_2_O_2_ production rates at the start (con) and 20 min after oxidation of substrates of reverse (RET) or forward (FET) electron transfer by intact mitochondria. **g,** Assessment of complex I activity before (con) and after 20 min of oxidation of various substrates. Aliquots were taken during incubation with substrates and NADH:HAR activity of complex I was measured. **h-i,** Representative flavin fluorescence scan of mitochondrial membrane complexes separated by hrCN PAGE and quantification of complex I-bound FMN in samples obtained after 20 min incubation with different substrates. Results are shown as mean ± s.d., n=4-12, *p<0.05, **p <0.01, ***p<0.001. Statistical significance was assessed with one-way ANOVA with Dunnett’s multiple-comparisons test (**e**, **i)** or t-test (**f**, **g**). Details of experiments are provided in Materials and Methods.

Previously, we showed that inhibition of complex I during RET is not due to disintegration of the hydrophilic head (N-module) of the enzyme^19^. Nevertheless, increased ROS generation from FMN of complex I during RET may contribute to oxidative damage of protein thiols^45^. To determine whether cysteine oxidation could cause the inhibition of mitochondrial complex I, we used the biotinylated iodoacetamide (BIAM) switch assay to analyze membranes before and after RET. The results failed to identify oxidized cysteine residues in the vicinity of FMN-binding site of complex I (*i.e.*, in NDUFV1, NDUFS1, NDUFV3, NDUFV2 subunits of the N-module). Moreover, most of the other identified mitochondrial proteins were found in a thiol-reduced state, except for several components of the thioredoxin system, which showed an increased abundance of oxidized thiols following RET (Extended Data Fig. 4 and Supplementary File 3). Together, these observations likely reflect both higher levels of NAD(P) reduction and increased ROS production within the mitochondrial matrix during RET.

### Characterization of AOX-expressing mouse

Prerequisite conditions for mitochondrial RET are a high ubiquinol/ubiquinone ratio in the inner mitochondrial membrane, and the presence of a sufficient proton-motive force to drive reverse electron flow upstream toward complex I. These are the exact metabolic conditions at the end of an oxygen deprivation period, where succinate and glycerol 3-phosphate oxidation promote ubiquinol accumulation, and the mitochondrial membrane becomes hyperpolarized due to a deficit of ADP for ATP-synthase (Fig. 2c-f, 2k,l). To manipulate mitochondrial electron flux during IR *in vivo*, we used transgenic mice that ectopically express alternative quinol oxidase (AOX) from *Ciona intestinalis*^46^. The presence of AOX creates a chimeric respiratory chain where the ubiquinol pool can be rapidly oxidized without generation of proton-motive force, as AOX bypasses complexes III and IV and transfers electrons from ubiquinol directly to molecular oxygen, eliminating the main prerequisite conditions that drive RET. Due to its low affinity for ubiquinol^9,47^, AOX is engaged only when the ubiquinone pool is highly reduced during RET (QH_2_/Q ratio is elevated). Therefore, we hypothesized that AOX presence would decrease RET and prevent over-reduction of complex I.

Hemizygous AOX mice develop normally and do not show any physiological abnormalities^48,49^. No significant differences were observed in the development of the cerebral vasculature between wild-type control and AOX-expressing mice (Fig. 5a, left). The expression of AOX was verified by genotyping, immunohistochemistry (Fig. 5a right), and western blotting (Fig. 5b). Confocal images of AOX staining (Fig. 5a right) show that AOX is mostly localized in the cellular cytoplasm with a punctate staining pattern indicating mitochondrial localization. Isolated brain mitochondria from AOX mice were found to catalyze direct oxidation of quinol by oxygen, and therefore respiration was only partially inhibited by the complex IV inhibitor cyanide (Fig. 5c). At the same time, the presence of AOX did not significantly affect state 2 and state 3 respiration (Fig. 5d). When respiring using RET substrates, AOX did not affect the efficiency of ATP-synthesis (Fig. 5e). Mitochondrial complex I content (as either an individual enzyme or as part its cognate of supercomplex), as well as total intramitochondrial NAD(P)H content (Fig. 5h-j) were not different in isolated mitochondria from control versus AOX mice. However, the degree of nucleotide reduction after the addition of RET substrate was lower in AOX-containing mitochondria compared to that in the control mice, indicating diminished RET-flow upstream of complex I (Fig. 5k-l). Importantly, ROS generation by AOX-containing intact mitochondria catalyzing RET was approximately half of the control activity (Fig. 5f,g). Overall, this indicates that AOX significantly decreases complex FMN reduction during RET. This conclusion was also supported by slower rates of ROS generation during succinate/glycerol 3-supported RET in mitochondria from AOX mice (Fig. 5m). Moreover, AOX expression attenuated RET-induced complex I inactivation (Fig. 5n) and prevented dissociation of FMN from the enzyme (Fig. 5o,p). Of note, the activity of matrix FMN phosphatase^43^ was not different between control and AOX-expressing mice (Extended Data Fig. 5a).

**Figure 5.**
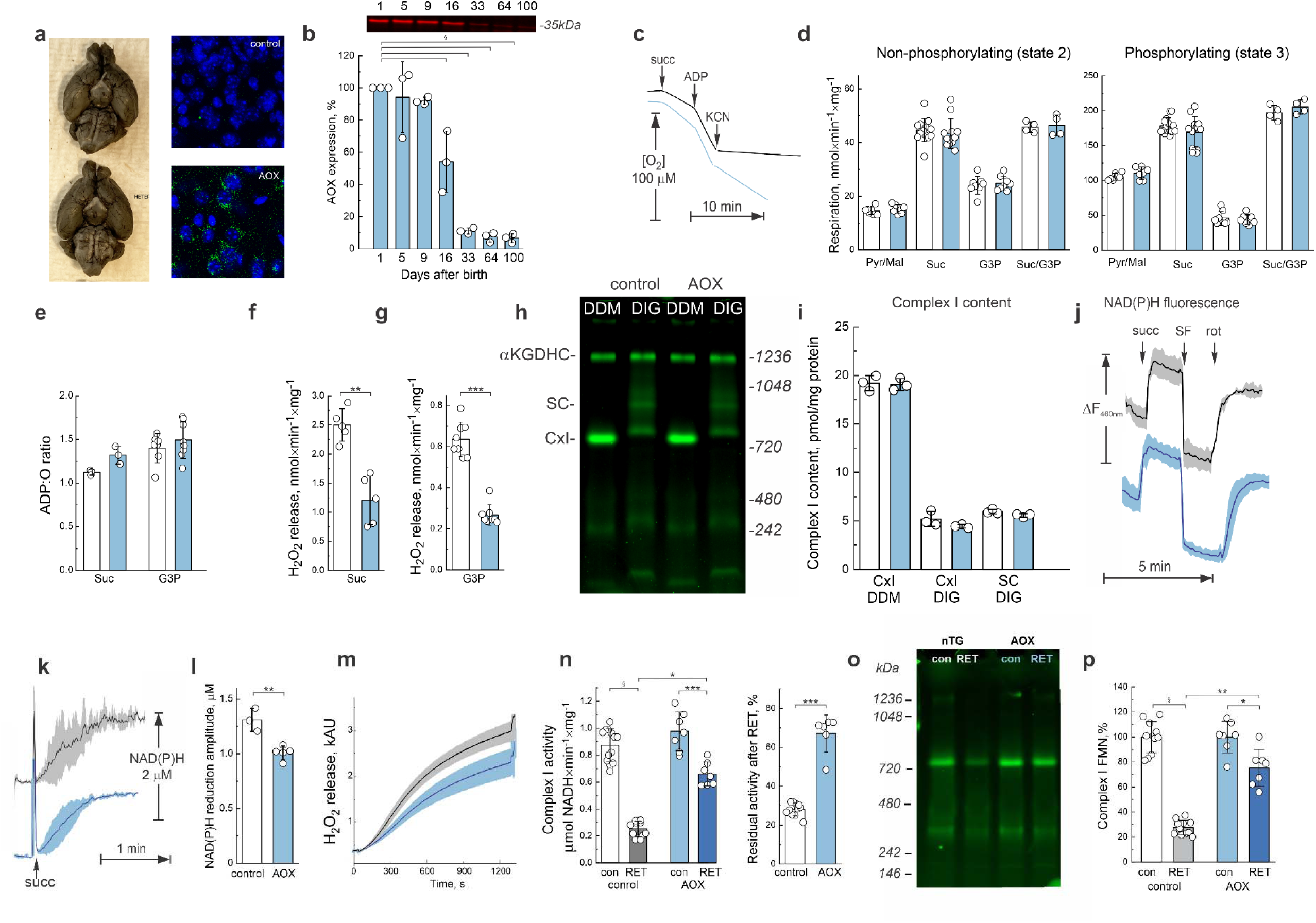
AOX expression attenuates RET-induced complex I inactivation. **a,** Left, images of control and AOX mouse brains with ink-perfused cerebrovasculature. Right, representative confocal images of immunofluorescence staining of brain cortex section from control and AOX-expressing mouse. Green, AOX-specific antibody and blue, DAPI staining. **b,** Representative western blotting image with antiserum against AOX using brain mitochondria from AOX-expressing mice at different ages. 30 µg of mitochondrial protein was applied to each well and signal was normalized by the intensity at 1 day of age. Quantification from three independent experiments. **c,** Representative trace of succinate-supported respiration of intact brain mitochondria from control (black) and AOX-containing (blue) mice. Reaction was started by the addition of 5 mM succinate to a standard media containing mitochondria (0.1 mg protein/ml). 0.5 mM ADP was added to initiate state 3 respiration, followed by the addition of 1 mM cyanide (KCN) as described in the Materials and Methods. **d,** Effect of AOX expression on state 2 (left) and state 3 (right) respiration of intact brain mitochondria isolated from control and AOX-containing mice (white and blue bars, respectively). Four types of substrates were used: a combination of malate/pyruvate, succinate, glycerol 3-phosphate and succinate/glycerol 3-phosphate. Reaction started by the addition of substrates and measured at 37°C as described in the Materials and Methods. **e,** Calculation of ADP:O ratio for control and AOX mice respiring on either succinate of glycerol 3-phosphate. **f-g,** ROS generation during oxidizing succinate (**f**) or glycerol 3-phosphate, (**g**) by control or AOX-containing mitochondria (white and blue bars, respectively). **h,** Representative flavin fluorescence scan of mitochondrial membrane complexes separated by rhCN PAGE and **i,** quantification of complex I-bound FMN content in samples from control or AOX-containing mitochondria (white and blue bars, respectively) solubilized with DDM (individual band of complex I) or digitonin (complex I is separated as individual enzyme (CxI) and as a part of supercomplexes (SC)). **j-l,** RET-induced NAD(P)H redox state in the matrix of control and AOX mitochondria. NAD(P)H autofluorescence was measured as described in Materials and Methods section. **j**, NADH(P) autofluorescence response upon addition succinate, uncoupler SF-6847 and complex I inhibitor rotenone. **k,** Time-resolved kinetics of RET-induced NAD(P)H reduction. Error bars represent one standard deviation based on the measurements of at least three separate mitochondria isolations. **l,** Quantification of amplitude of NAD(P)H reduction. **m,** Decline of H_2_O_2_ generation by intact mitochondria during RET supported by oxidation of succinate/glycerol 3-phosphate. Error bars represent one standard deviation based on triplicate measurements of different mitochondria isolations. **n,** Assessment of complex I activity before (con) and after 20 min of oxidation of various substrates. Aliquots were taken during incubation with substrates and NADH:HAR activity of complex I was measured. Comparison of absolute values (left) and fractions of residual activity after RET (right). **o,** Representative flavin fluorescence scan of mitochondrial membrane complexes separated by rhCN PAGE and **p,** quantification of complex I-bound FMN in samples obtained before (con) and after 20 min incubation with a combination of succinate/glycerol 3 phosphate (RET) with different substrates. In all panels data for control or AOX-containing mitochondria are shown in white or blue, respectively. Values are shown as mean ± SD, n=4-12. Statistical significance was assessed with one-way ANOVA with Dunnett’s multiple comparisons **(b),** two-way ANOVA with Tukey’s post-hoc (**n**,**p**) or t-test (**d**,**e**-**g**, **i**, **I**,**o**).

### AOX expression is neuroprotective in models of IR in vitro and in vivo

We examined how the expression of AOX modulated hippocampal injury caused by oxygen-glucose deprivation (OGD) followed by reoxygenation. Hippocampal slices prepared from control animals subjected to OGD showed substantial injury in the CA1 region and cell death. The expression of AOX reduced the size of the injury, establishing a neuroprotective effect of AOX in IR (Fig. 6a,b and Extended data Fig. 6).

**Figure 6.**
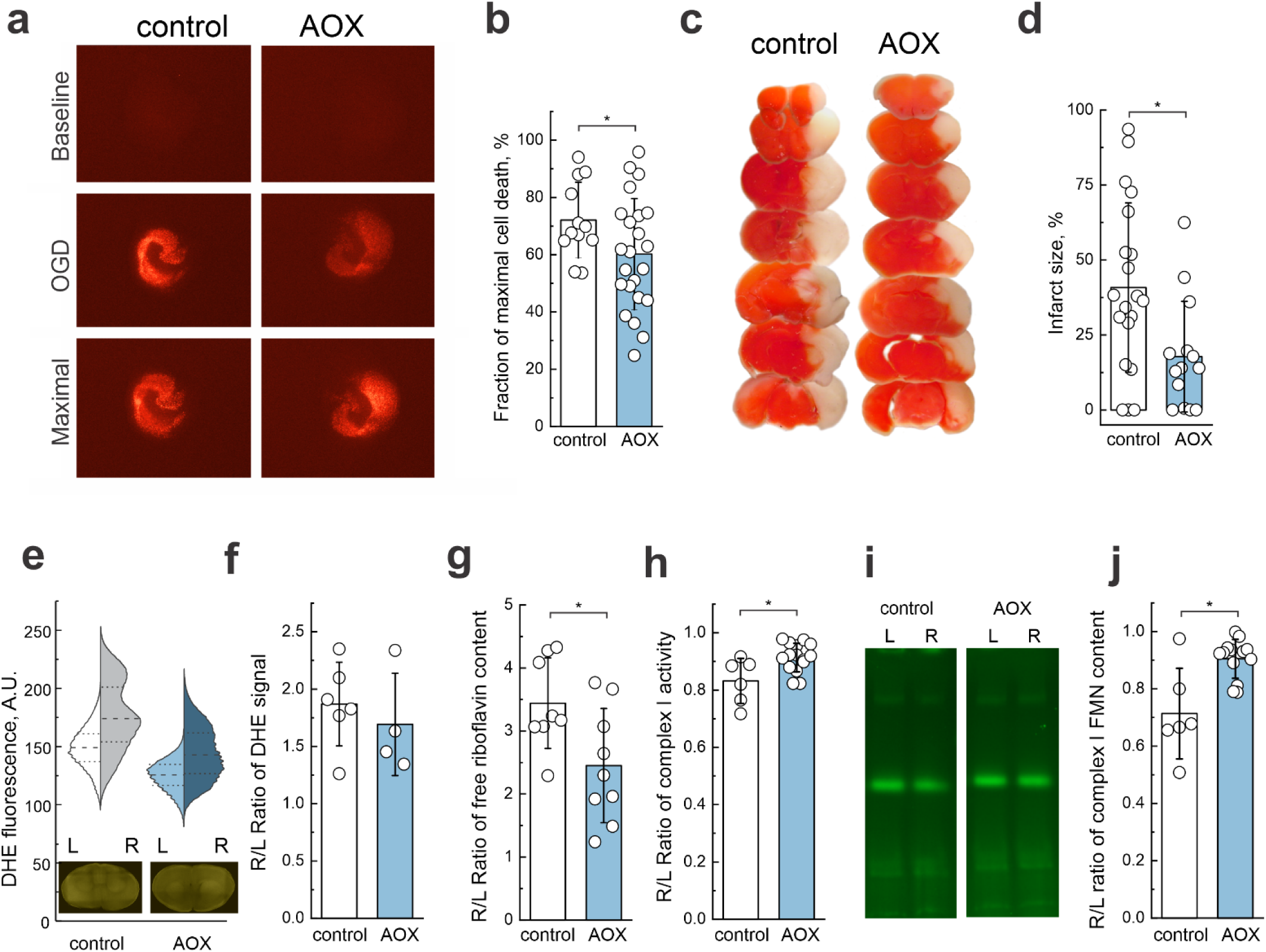
AOX expression reduces ischemic damage and protects mitochondrial function. **a,** Representative live/dead staining images of brain slices from control and AOX-expressing mice under baseline conditions, after OGD, and at maximal cell death. **b,** Quantification of cell death expressed as a fraction of maximal cell death, showing reduced susceptibility in AOX mice. **c,** TTC-staining of coronal brain sections 24 h after HI-reperfusion. Functional tissue was stained (red), and the infarcted tissue was not stained (white). **d,** Infarct volume is shown as a percentage of the hemisphere volume, demonstrating reduced infarct area in AOX mice. **e,** Violin plots of dihydroethidium (DHE) fluorescence intensity in the left and right hemispheres, with corresponding representative images of brain section from control and AOX mice after HI-reperfusion (inset). **f,** Quantification of DHE signal ratios from the right/left hemispheres, showing no significant differences between genotypes. **g,** Ratio of free riboflavin content (right/left hemispheres), measured in samples after HI in control and AOX mice. **h,** Ratio of mitochondrial complex I activity (between right and left hemispheres), showing enhanced activity in AOX mice after IR. **i,** Representative flavin fluorescence scan of hrCN PAGE of mitochondrial complexes from samples obtained from left and right hemispheres of control and AOX mice after HI-reperfusion *in vivo*. **j,** Ratio of complex I-bound FMN content ratio between right and left hemispheres, showing increased FMN content in AOX mice. All data are presented as mean ± s.d., with individual data points shown. Statistical significance was determined using t-tests.

Using an *in vivo* model of brain IR we then measured the effect of AOX expression on infarct size. Notably, after 24 h of reperfusion, mice expressing AOX exhibited 50% lower cerebral infarct volumes compared to control WT littermates (Fig. 6c,d). Next, we assessed the level of ROS in the ipsi- and contralateral hemispheres 4 h after reperfusion by dihydroethidium (DHE) labeling. Upon *ex vivo* DHE fluorescence imaging of brain samples, we observed a significantly higher ROS level in the ipsilateral right hemisphere for both genotypes (Fig. 6e, Extended Data Fig. 7). However, when the ratio of total DHE fluorescence signals between ipsi- and contralateral hemispheres was measured, we did not detect a significant difference between AOX and control mice (Fig. 6f). Metabolomics studies (Supplementary Table 4) showed no significant difference in the levels of succinate and glycerol 3-phosphate in the brains of control and AOX mice after oxygen deprivation. However, the level of free riboflavin was significantly lower in the ipsilateral area of the brain in AOX-expressing mice when compared with the control (Fig. 6g). Importantly, complex I activity and enzyme-bound FMN content were higher in mitochondria from AOX expressing mice (Fig. 6h-j). Taken together, these findings indicate that AOX does not only significantly decrease IR-induced ROS generation, but also prevents the dissociation of FMN from mitochondrial complex I. Collectively, our findings demonstrate that AOX expression confers neuroprotection by mitigating ischemia-induced cell death and decreasing infarct size.

## Discussion

Overall, our study provides an unprecedented longitudinal analysis of metabolic flux changes following mouse brain IR, elucidating the mechanisms underlying mitochondrial failure. We demonstrate that IR-induced complex I inactivation occurs after flavin dissociation, driven by RET. Using AOX-expressing mice, which exhibit lower rates of RET, we were able to mitigate FMN loss from mitochondrial complex I after brain IR resulting in neuroprotection (Fig. 7).

**Figure 7.**
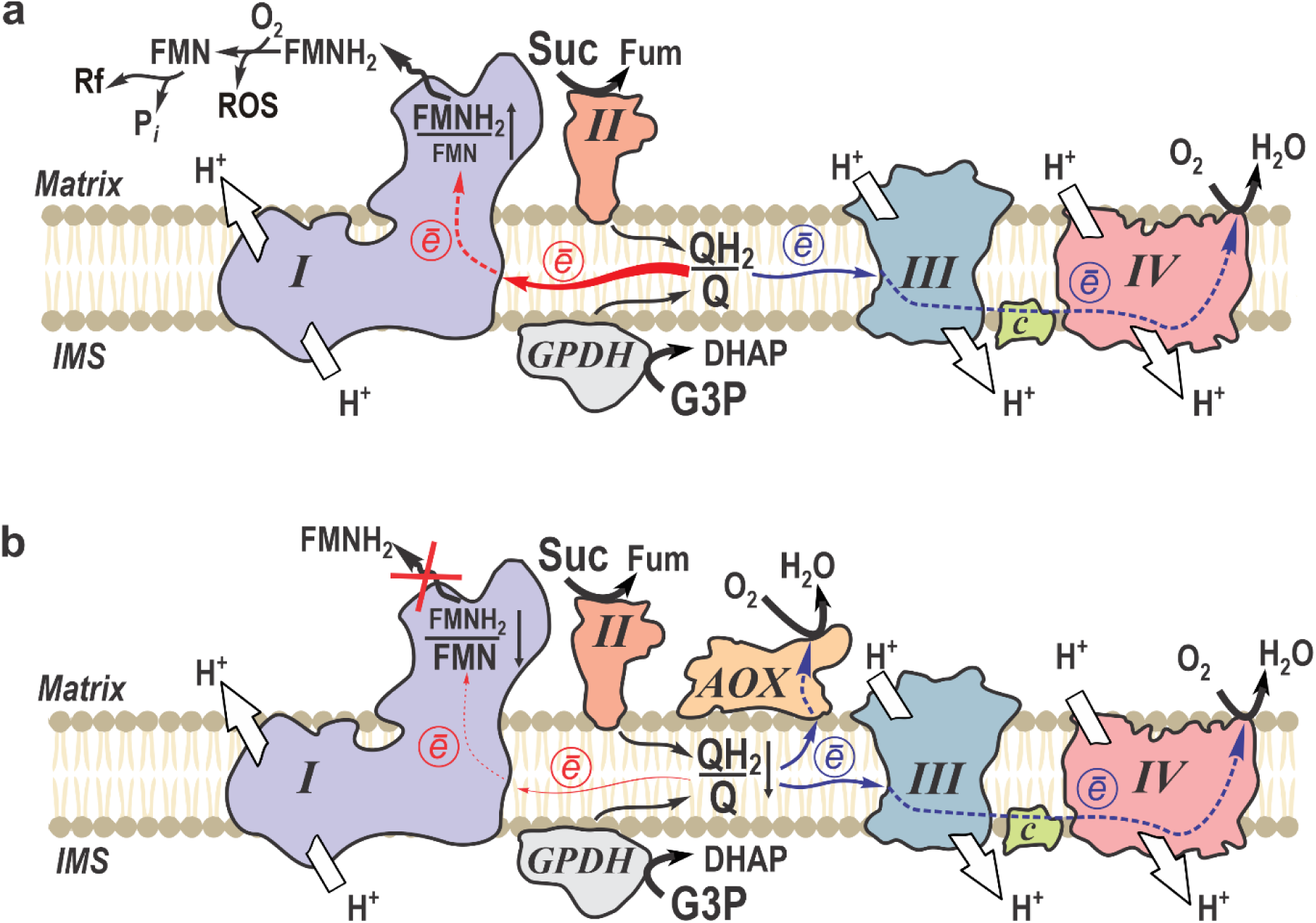
Respiratory chain rewiring in mitochondria of AOX-expressing mice during early reperfusion phase in brain IR. **a**, Upon reperfusion mitochondria with canonical respiratory chain oxidize RET-substrates as succinate (Suc) and glycerol 3-phospahe (G3P) accumulated during ischemic phase. Complex III (III) and cytochrome *c* oxidase (IV) function normally in the forward electron transfer mode (e^-^, blue arrows), pumping protons across the inner mitochondrial membrane from the matrix side to the intermembrane space (IMS) to sustain proton-motive force. Rapid reduction of ubiquinone pool (high QH_₂_/Q ratio) supports RET from complex II (II) and glycerol 3-phosphate dehydrogenase (GPDH) towards complex I (I) (RET is shown by red arrows). Overreduction of complex I leads to FMNH_₂_ dissociation, followed by autooxidation by molecular oxygen with non-enzymatic ROS formation and degradation of FMN to riboflavin (Rf). **b,** Expression of AOX in mitochondria affords redirection of electrons from quinol (QH_2_) to oxygen without proton pumping, bypassing complexes III and IV. This bypass limits RET toward complex I, therefore attenuating dissociation of the reduced flavin and protecting complex I from inactivation.

Our findings identify the mechanism of RET-induced complex I inactivation: excessive enzyme reduction, leading to the dissociation of FMN from the enzyme. RET is fueled by the oxidation of succinate and glycerol 3-phosphate, which accumulated during oxygen deprivation. The buildup of succinate during ischemia in the brain and other organs has been described before^26,39,50-53^, but here we report a dramatic increase of glycerol 3-phosphate as a major RET-supporting substrate^8^. We also found degradation of nearly half of the adenylate pool at the time of reperfusion and therefore propose that a shortage of ADP limits the activity of mitochondrial F_1_F_o_ ATP synthase. This abates consumption of the proton-motive force, which in turn promotes RET, exacerbating mitochondrial dysfunction. After the complete reduction of matrix nucleotides in the QH_2_:NAD^+^ reductase reaction^21,22^, steady-state RET-driven electron pressure maintains redox centers of complex I, including FMN, in the over-reduced state. This promotes the escape of electrons to oxygen with dramatic ROS generation upstream of the quinone binding site of complex I^16,17,23,35,37,38,52,54^. Importantly, the reduced form of flavin exhibits lower affinity for its binding site in complex I, leading to its slow dissociation, with a reported *in vitro* half-time of several minutes^17,55-57^. The resulting FMN-deficient complex I is unable to catalyze both its physiological reaction of oxidation of NADH by quinone or the production of ROS. *In vivo*, liberated reduced FMN undergoes autoxidation by molecular oxygen, since even during HI the residual oxygen level is estimated at 4-5 µM (Fig. 1). FMN is then metabolized by matrix phosphatase^42,43^, leading to a dramatic increase of free riboflavin in the tissue after oxygen deprivation. Lack of ATP during the early reoxygenation phase may slow resynthesis of FMN by riboflavin kinase and, therefore, further delay the reactivation of complex I.

Our data identify FMN dissociation from complex I as a key event in mitochondrial failure during brain IR injury. This mechanism, previously unrecognized in the context of neurodegeneration, redefines how mitochondrial impairment drives tissue damage. Importantly, its translational significance is underscored by our recent clinical work showing that FMN release from complex I is a decisive marker of liver graft quality prior to transplantation ^58^ ^59^. Together, these findings position FMN dissociation not only as a fundamental driver of IR pathology but also as a promising therapeutic and diagnostic target across organs.

Expression of AOX, which substitutes for the activities of complexes III and IV except for their proton translocating activity, results in oxidation of ubiquinol produced by succinate dehydrogenase or GPDH during IR (Fig. 7b). In turn, this decreases the level of RET in complex I, evident from lower RET-induced ROS release and matrix NADH reduction. Previously, a decrease of ROS production in the presence of AOX has been shown *in vitro* for isolated plant^60^ and mammalian mitochondria^9,49,61^. This study provides the first evidence that AOX expression mitigates RET, preserves complex I FMN binding, and promotes neuroprotection, suggesting that preventing complex I reductive inactivation may be beneficial in IR brain injury.

Mitochondrial electron flux via RET and escalation of ROS generation are recognized as key steps in the development of brain IR injury^16,23,25,26,52^. We detected higher ROS production in the affected hemisphere after IR, but surprisingly no significant difference was found between control and AOX-expressing mice, although AOX-containing mitochondria generate only half the level of ROS on RET-supporting substrates *in vitro*. Our findings call for a re-evaluation of ROS sources in brain IR. Contrary to the prevailing view that mitochondrial ROS are the primary drivers, our *in vivo* data suggest that RET may not be the main contributor to the IR-induced ROS burst. Instead, AOX confers neuroprotection by diverting electrons from the quinone pool to oxygen and preventing overreduction and FMN dissociation from complex I.

In summary, our study uncovered a novel mechanism of mitochondrial complex I inactivation driven by RET and FMN dissociation during the early phase of brain IR. By identifying the metabolic and redox conditions that promote RET, such as substrate accumulation and adenylate depletion, we clarified how mitochondrial dysfunction is initiated and sustained in IR injury. Crucially, the use of AOX highlights a viable strategy to bypass RET-induced stress by rewiring electron transport in the respiratory chain and preserving complex I integrity. These findings not only provide new mechanistic insight into mitochondrial failure in brain IR but also support AOX-based interventions as a promising avenue for neuroprotection in IR brain injury.

### Limitations of this study

Our findings identify RET-induced FMN dissociation as a critical mechanism of complex I inactivation and demonstrate the neuroprotective effect of AOX *in vivo*. Further work will be required to investigate the extent to which this phenomenon occurs in different models of IR brain injury. While AOX expression is not currently translatable to clinical practice, the neuroprotection observed in this study— as well as with electron mediators such as methylene blue^62^ – highlights the therapeutic potential of respiratory chain electron flux rewiring as a compelling strategy for intervention during the acute phase of brain IR. Although AOX expression effectively attenuates RET and ROS production in isolated mitochondria, the *in vivo* reduction in ROS was less pronounced, suggesting that non-mitochondrial sources may contribute to IR-induced oxidative stress. Notably, AOX’s ROS-diminishing capacity increases with decreasing temperature^61^, implying that its efficacy may be synergetic with hypothermic treatment. Additionally, the use of a neonatal mouse model may limit direct extrapolation to adult or human ischemic conditions. Finally, while we showed a correlation between AOX-mediated complex I protection and neuroprotection, the precise downstream signaling pathways and long-term functional outcomes warrant further investigation.

## Methods

### Reagents

A complete list of the reagents used in this study is included in Supplementary Table 5.

### Mice

Mice were used at 10 days of age (P10). All animal studies were conducted according to Federal and University regulations and protocols approved by Weill Cornell Medicine and by the Columbia University Institutional Animal Care and Use Committees (IACUC). Wild-type C57BL/6J mice were obtained from Jackson Laboratories (strain #000664) and bred in-house at Weill Cornell Medicine. We used a previously generated transgenic mouse line, which contains a lox-stop-lox-AOX targeting construct in the Rosa26 locus^46^. Hemizygous mice expressing AOX protein in all tissues^48,61^ (referred to as AOX mice) were generated by breeding C57BL/6Rosa26^AOX-lsl/wt^ with B6.FVB-*Tmem163*^Tg(ACTB-cre)2Mrt^/EmsJ mice (The Jackson Laboratories, strain #019099). Littermates not expressing Cre-recombinase were used as wild-type controls (control mice). Hemizygous AOX mice exhibited normal development, with no observable phenotypic abnormalities.

### Neonatal cerebral hypoxia-ischemia (HI) injury

Transient hypoxia-ischemia (HI) was induced as described^29^. HI brain injury was induced on P10 mice by permanent ligation of the right common carotid artery under isoflurane anesthesia. Following 1.5 h of post-surgical recovery, mice were subjected to hypoxia (8% O_2_/ 92% N_2_, Tech Air Inc., NY) for 15 min, at 37 ± 0.5°C. This model mimics IR brain injury caused by neonatal stroke and birth asphyxia (global reperfusion due to acute placental failure at birth or hypoxic cardiac arrest after birth up to 28 days of life). This model of HI-reperfusion is highly characteristic of perinatal IR injury, and benefits from quantifiable measures of brain infarction in the ipsilateral (right) hemisphere leading to permanent neurofunctional deficit^28,29^.

Immediately after HI or after a period of recirculation (15 min, 1, 2, 4, 6, or 24 h), mice were decapitated, and heads were rapidly frozen in liquid nitrogen for further mitochondrial membranes preparation. At 24 h of reperfusion, another cohort of animals was used for assessment of the extent of cerebral injury.

### Measurement of infarct volume

At 24 h of reperfusion mice were euthanized, fresh brains were collected, sectioned into 1 mm thick coronal slices and stained with 2% triphenyltetrazolium chloride (TTC). Digital images of infarcted (white, no staining) and viable (red) areas of brains were traced (Adobe Photoshop 4.0.1) and analyzed (NIH image 1.62J). The extent of brain injury was expressed as a percentage of the right hemisphere ipsilateral to the carotid artery ligation side.

### ROS imaging in *vivo*

Intraperitoneal injection of fresh dihydroethidium (DHE, 5 mg/kg) was administered to 10-day-old mice, 5 minutes prior to permanent ligation of the right common carotid artery. IR *in vivo* protocol was conducted as described above. After 6h reperfusion mice were decapitated, and the brain was quickly removed. Next, coronal slices were performed (1 mm-thick) and placed onto glass slides. The fluorescence produced from DHE oxidation in each slice was measured using ImageXpress Pico System with the TRITC filter cube (Excitation/Emission = 545/594 nm). Fluorescence intensity in the regions of interest (ipsi and contralateral hemispheres) was analyzed as arbitrary units with background correction applied using ImageJ, napari, and OriginPro (version 9.8) software. The total fluorescence for each hemisphere in the brain was calculated by adding together the fluorescence values from all individual slices and shown as a violin plot. Additionally, a comparison of fluorescence was conducted by dividing the signal intensity in the right hemisphere by that in the left hemisphere.

### Oxygen measurements *in situ*

We monitored brain oxygen level in the affected hemisphere by direct, non-invasive measurements of tissue oxygen with the phosphorescence quenching method^63^ using oxygen probe Oxyphor PdG4^30^. Oxygen levels were derived from phosphorescence decay lifetimes, which are independent of the local probe concentration and/or optical properties (*e.g.*, absorption and scattering heterogeneities) of the environment. OxyphorPdG4 was injected into the jugular vein during surgery to achieve ∼1 µM final concentration in the blood plasma. The measurements were performed using a fiberoptic phosphorometer (Oxyled, Oxygen Enterprises) equipped with a laser diode for excitation (λ_ex_=635 nm). The laser was focused into a ∼200 µm spot in the area corresponding to the expected infarction (2 mm back from the bregma and 3.5 mm to the right), and the detection fiber (4 mm diameter) was positioned next to the excitation focus. In this setup, the phosphorescence probes tissue oxygenation up to 2.5 mm below the skin level. Phosphorescence decay lifetime was converted to oxygen concentration using a Stern-Volmer-like calibration plot measured independently (Extended Data Fig. 1)

### Oxygen-glucose deprivation (OGD) and reperfusion in hippocampal slices

Hippocampal slice cultures (coronal slices, 350 μm thick) were prepared as previously described^64^ and cultured on Millicel CM membrane inserts (Millipore). After 14 days in culture, slices were imaged for three consecutive days after staining with propidium iodide (PI) in the culture medium (5 μg/mL). On the first day, baseline PI images were captured (Fl_basal_) in a transmission fluorescence microscope with a 2× objective, and then OGD was performed. Briefly, slices were washed in OGD buffer (125 mM NaCl, 5 mM KCl, 1.2 mM Na2PO4, 26 mM NaHCO3, 1.8 mM CaCl2, 0.9 mM MgCl2, 10 mM HEPES, pH 7.4), and then incubated in OGD buffer without glucose in an anoxic gas chamber for 1 h, before they were returned to normal culture medium. On the second day, post-OGD PI images were captured (Fl_OGD_). Then, slices were treated with 1 mM N-Methyl-D-aspartic acid (NMDA) for 24 h. On the third day, post-NMDA PI images were captured, representing the maximum death in each slice (Fl_max_). The same imaging settings were used at all time points, and images from the CA1 hippocampal region were used to calculate the proportion of OGD-induced death using the formula (Fl_OGD_ − Fl_basa_l)/(Fl_max_ − Fl_basal_) × 100%.

### Immunohistochemical staining in coronal brain slices

AOX transgenic mice and littermate control wild-type mice (8-10 days of age) were anesthetized in CO_2_ and decapitated. The brains were removed and placed into 4% paraformaldehyde solution in phosphate buffer (pH 7.4) overnight at 4°C on a shaker. The following day, the brains were mounted, and slices were cut 40 μm in thickness with a vibratome. The brain slices were permeabilized in 0.25% Triton X-100 in PBS for 1 h followed by incubation in 10% normal donkey serum in PBS for 2 hrs. The slices were then incubated with diluted AOX primary antibody (21st Century Biochemicals 1:100) for 72 h in a cold room with shaking. The slices were washed 3 times with PBS and placed into a solution containing donkey anti-rabbit secondary antibody conjugated to Alexa 488 for 2 h at room temperature. The slices were then incubated with DAPI solution (0.5 μg/ml) for 20 min. After washing 3 times in PBS, the slices were mounted and viewed with a confocal microscope (Leica SP8). The images were captured with a 20x optical lens and magnified 3x with digital zoom. The settings remained the same during imaging acquisition for both AOX and control samples.

### Preparation of brain mitochondrial membranes for ex vivo studies

Mitochondrial membranes were isolated from frozen ipsilateral brain hemispheres by differential centrifugation. After sagittal transection of the frozen heads into two hemispheres, the regions with maximal damage were excised from the ipsilateral hemisphere cortex caudal to bregma level. Pieces of frozen brain tissue were homogenized with 60 strokes of tight pestle of 2 ml Kontes™ Dounce homogenizer in 1 ml of ice-cold isolation buffer MSE (225 mM mannitol, 75 mM sucrose, 20 mM HEPES-Tris, 1mM EGTA, 1mg/ml BSA, pH 7.4), containing 80 µg/ml alamethicin to release low molecular weight metabolites from mitochondria. Tissue debris was discarded after the centrifugation at 1,500g for 5 min at 4□°C. The supernatant was centrifuged at 20,000g for 15 min at 4□°C, and the membrane pellet was washed twice with the isolation buffer without BSA. The final pellet was resuspended in 60 µl of the same buffer and stored at -80□°C until use.

### Isolation of intact brain mitochondria for in vitro studies

Intact brain mitochondria were isolated from neonatal or adult mice by differential centrifugation with digitonin treatment^17^. Forebrain hemispheres were excised and immediately immersed into ice-cold MSE buffer. One brain was homogenized with 40 strokes by tight pestle of a Dounce homogenizer in 10 ml of the MSE buffer, diluted twofold and centrifuged at 5,000 g for 4 min at 4□°C in a refrigerated Beckman centrifuge. The supernatant was supplemented with 0.02% digitonin and centrifuged again at 10,000 g for 10 min. The pellet was then washed twice by centrifuging at 10,000 g for 10 min in MSE buffer without BSA. The final pellet was resuspended in 0.15 ml of washing buffer, supplemented with 1 mg/ml BSA, and stored on ice. At least three separate isolations were used for each experimental condition.

### Respiration and H_2_O_2_ measurements in intact mitochondria

Mitochondrial respiration and extramitochondrial release of H_2_O_2_ were measured using a high-resolution respirometer (O2k oxygraph, Oroboros Instruments, Innsbruck, Austria) equipped with two-channel fluorescence optical setup to monitor simultaneously oxygen level and fluorescence (excitation /emission 525/580 nm)^8^. Mitochondria (0.1 mg of protein) were added to 2 ml respiration buffer composed of 125 mM KCl, 0.2 mM EGTA, 20 mM HEPES-Tris, 4 mM KH_2_PO_4_, pH 7.4, 2 mM MgCl_2_, 1 mg/ml BSA, 10μM Amplex UltraRed (Invitrogen), 4 U/ml horseradish peroxidase, and 5 U/ml superoxide dismutase at 37°C. The following substrates were used: 2 mM malate and 5 mM pyruvate (for complex I-supported respiration), 5 mM succinate (for complex II) and 1 mM glutamate, or 40 mM glycerol 3-phosphate (for GPDH). After recording oxygen consumption in non-phosphorylating conditions, 200-500 μM ADP was added to initiate the State 3 (phosphorylating) respiration. Respiration was fully sensitive (inhibited) by 1 mM cyanide or 1 μM antimycin A. The H_2_O_2_ production was followed by the raw resorufin fluorescence and calibrated at the end of the run by adding 200-400 pmoles of a fresh standard solution of H_2_O_2_ (ε_240nm_ = 46.3 M^-1^ cm^-1^) to the chamber containing all the components of the assay.

### Measurement of respiratory chain enzymes activities in mitochondrial membranes

All activities were measured spectrophotometrically using Molecular Devices SpectraMax M5 plate reader in 0.2 ml of the assay buffer (125 mM KCl, 20 mM HEPES-Tris, 0.02 mM EGTA, pH 7.4) at 25□°C.

NADH-dependent activities of complex I were assayed as oxidation of 0.15 mM NADH at 340 nm (ε_340nm_ = 6.22 mM^-1^cm^-1^) in the assay buffer supplemented with 40 µg/ml alamethicin, and 1 mM KCN (NADH media). NADH:Q reductase was measured in NADH media containing 1 mg/ml BSA, 60 µM decylubiquinone, and 5-15 µg protein per well. Only rotenone (1 µM) sensitive part of the activity was used for the calculations. NADH:HAR reductase was assayed in NADH media containing 1 mM HAR and 2-5 µg protein per well.

Complex II succinate:DCIP reductase activity was recorded at 600 nm (ε_600nm_ = 21 mM^-1^cm^-1^) in the assay buffer containing 5 mM succinate, 40 µM decylubiquinone, 0.1 mM 2,6-dichlorophenol-indophenol (DCPIP), 1 mM KCN, and 5-10 µg protein per well. Activity was fully sensitive to 5 nM atpenin, a specific complex II inhibitor.

Dihydroorotate dehydrogenase (DHODH) activity was determined at 37°C as DCPIP reduction essentially as described elsewhere ^65^ at 600 nm (ε_600nm_ = 21 mM^-1^cm^-1^) in the assay buffer containing 0.1% Triton X-100, 10 mM DHO, 40 µM decylubiquinone, 50µM DCIP, 2 mM KCN, 2 µM rotenone and 1 µM antimycin, and 20-40 µg protein per well. Reaction was 90% sensitive to the addition of 0.4 µM brequinar.

Complex IV oxidase activity was measured as oxidation of 50 µM ferrocytochrome *c* at 550 nm (ε_550 nm_ = 21.5 mM^-1^cm^-1^) in the assay buffer supplemented with 0.025% N-dodecyl-β-D-maltoside and 1-3 µg protein per well. The activity was fully sensitive to 0.5 mM cyanide.

FMN phosphatase activity was measured in preparation of matrix fraction of intact mitochondria in the assay buffer. First, 0.4 mg/ml mitochondria were preincubated with 40 µg/ml alamethicin for permeabilization, and 100 µM of FMN was added. Aliquots were taken in time and concentration of inorganic phosphate was measured using Malachite green assay (Sigma #MAK307). Increase in free phosphate was linear up to 40 µM level within the first 15 min of reaction.

### Mitochondrial NADH autofluorescence

Matrix NAD(P)H autofluorescence in mitochondria oxidizing different substrates was measured by Hitachi 7000 spectrofluorimeter set at excitation/emission = 350/460 nm^21^. The reaction was carried out under the same conditions as the respiratory assays with 0.2 mg/ml mitochondrial protein in 300 µl of the assay buffer. Amplex UltraRed and horseradish peroxidase were omitted from the assay. Additions to the suspensions were succinate (5 mM), SF (35 nM), rotenone (1 µM). Special controls for photobleaching of the preparation during prolonged measurements were performed. Signal was calibrated by the addition of aliquots of standard NADH solution at the end of the experiment.

### Complex I FMN content measurements

Sample solubilization with DDM and running conditions for high-resolution clear native (hrCN) electrophoresis of mitochondrial membranes and tissues were performed as previously described^44,66^. Gels were scanned for flavin fluorescence in a Typhoon 9000 gel scanner (GE) using a 473 nm laser and BPB1 filter (530 nm maximum, 20 nm bandpass). Flavin signal calibration was performed by adding standard solutions of FMN (1-10 pmoles) directly onto the gel.

### AOX expression detection by western blotting

Whole-tissue homogenates were obtained as described above, and protein concentration was determined by the Pierce bicinchoninic acid assay (ThermoFisher A55865). Homogenate (20 μg protein) fractions were denatured in 1X Laemmli Buffer (Bio-Rad) containing 2-Mercaptoethanol at 95°C for 10 min and separated by electrophoresis in a 4–12% SDS–PAGE gel (Bio-Rad) and transferred to a PVDF membrane (Bio-Rad). Blots were incubated in 3% BSA in TBS with 1% Tween-20 (TBST) for 1 h at room temperature. Primary AOX antibodies (AOX, 21st Century Biochemicals, customized rabbit polyclonal, 1:50,000) were incubated overnight at 4□°C. Secondary antibodies (HRP-antirabbit, Jackson ImmunoResearch,1:10,000) were incubated for 45 min at room temperature in the dark. After washing all blots with TBST or TBS, the proteins were imaged on Odyssey Licor (Bio-Rad) as described previously^67^.

### BIAM switch redox proteomics and MS analysis

To assess the effect of RET on the thiol redox state *in vitro*, intact brain mitochondria from P10 mice were incubated in a medium 125 mM KCl, 0.2 mM EGTA, 20 mM HEPES-Tris, 4 mM KH_2_PO_4_, pH 7.4, 2 mM MgCl_2_, 1 mg/ml BSA with or without 10 mM succinate. After 20 min aliquots containing 35 µg of protein were taken, and cold 100% TCA was added to final concentration of 20% TCA and aliquots were frozen in liquid nitrogen.

Detailed description of redox proteomics, raw and analyzed data have been deposited to the ProteomeXchange Consortium (http://proteomecentral.proteomexchange.org) via the PRIDE partner repository^68^ with the dataset identifier PXD063555. For data analysis, MaxQuant v1.6.1.0 was used^69^. Proteins were identified using mouse reference proteome database UniProtKB with 52538 entries, released in 2/2018. Acetylation (+42.01) at N-terminus, NEM (+125.05), and BIAM (+414.19), and oxidation of methionine (+15.99) were selected as variable modifications. The enzyme specificity was set to Trypsin. False discovery rate (FDR) for the identification of proteins and peptides was 1%. The resulting oxidized peptide data was filtered to remove reverse hits and common contaminants. Any peptide identified in fewer than 4 of the 12 biological samples in at least one group was excluded from further analysis. Next, global sums of non-oxidized LFQ intensities were also compared by an unpaired t-test to assess overall differences in sample loading or total proteome changes. No significant difference was found between control and RET total protein. Missing intensities (zero LFQ values) were imputed from the lowest 5% of the observed distribution in each sample (i.e., random draws from the low-intensity tail. To compare control *vs.* RET conditions, we performed a paired t-test on the LFQ BIAM intensities. For visualization, a volcano plot was generated with the log_₂_(fold change) on the x_-_axis and the −log_₁₀_(p_-_value) on the y_-_axis. Proteins exceeding a fold-change of 2 at p < 0.05 were highlighted as significantly differentially oxidized. The python script used for this analysis is available in the Supplementary File 6.

### LC/MS metabolite extraction

Brain samples were extracted using -70°C 80:20 methanol:water. The tissue-methanol mixture was subjected to bead-beating for 45 sec using a Tissuelyser cell disrupter (Qiagen). Extracts were centrifuged for 5 min at 5,000 rpm to pellet insoluble protein, and supernatants were transferred to clean tubes. The extraction procedure was repeated two additional times, and all three supernatants were pooled, dried in a Vacufuge (Eppendorf) and stored at -80□°C until analysis. The methanol-insoluble protein pellet was solubilized in 0.2 M NaOH at 95□°C for 20 min, and protein was quantified using a BioRad DC assay. On the day of metabolite analysis, dried cell extracts were reconstituted in 70% acetonitrile at a relative protein concentration of 4 µg/ml, and 4 µl of this reconstituted extract was injected for LC/MS-based targeted and untargeted metabolite profiling.

### LC/MS metabolomics platform for untargeted metabolite profiling

Tissue extracts were analyzed by LC/MS as described previously^70^, using a platform comprised of an Agilent Model 1290 Infinity II liquid chromatography system coupled to an Agilent 6550 iFunnel time-of-flight MS analyzer. Chromatography of metabolites utilized aqueous normal phase (ANP) chromatography on a Diamond Hydride column (Microsolv). Mobile phases consisted of: (A) 50% isopropanol, containing 0.025% acetic acid, and (B) 90% acetonitrile containing 5 mM ammonium acetate. To eliminate the interference of metal ions on chromatographic peak integrity and electrospray ionization, EDTA was added to the mobile phase at a final concentration of 5 µM. The following gradient was applied: 0-1.0 min, 99% B; 1.0-15.0 min, to 20% B; 15.0-29.0, 0% B; 29.1-37min, 99% B. Raw data were analyzed using MassHunter Profinder 10.0 and MassProfiler Professional (MPP) 15.1 software (Agilent technologies). Student t-tests (p < 0.05) were performed to identify significant differences between groups.

### LC/MS Metabolite Structure Specification

To ascertain the identities of differentially expressed metabolites (p < 0.05), LC/MS data were searched against an in-house annotated personal metabolite database created using MassHunter PCDL manager 8.0 (Agilent Technologies), based on monoisotopic neutral masses (<5 ppm mass accuracy) and chromatographic retention times. A molecular formula generator (MFG) algorithm in MPP was used to generate and score empirical molecular formulae, based on a weighted consideration of monoisotopic mass accuracy, isotope abundance ratios, and spacing between isotope peaks. A tentative compound ID was assigned when PCDL database and MFG scores concurred for a given candidate molecule. Tentatively assigned molecules were verified based on a match of LC retention times and/or MS/MS fragmentation spectra for pure molecule standards contained in a growing in-house metabolite database.

### MALDI MS Sample preparation

Mouse heads were frozen in liquid nitrogen and stored at -80□°C until processing. Brain cryosections were cut at 10 µm thickness, mounted on conductive slides coated with indium tin oxide (ITO; Delta Technologies; cat # CB-90IN-S111) and stored at -80□°C. On the day of MALDI MS data acquisition, ITO-slides with tissue sections were transferred to a vacuum chamber and dried for 30 min prior to deposition of matrices for imaging: N-(1-naphthyl)ethylenediamine dihydrochloride (NEDC; Sigma Aldrich, cat # 1465-25-4) at 10 mg/ml in 75% methanol, and 9-aminoacridine (9AA, Sigma Aldrich, cat # 90-45-9) at 5mg/ml in 85% Ethanol, respectively. All matrices were delivered using an HTX TM-Sprayer™ (HTX Technologies LLC, NC) with optimized spraying parameters for each individual matrix. Matrix-coated tissue sections were dried in a vacuum desiccator for 20 min before MALDI MS imaging data acquisition in negative ion detection mode.

### MALDI MSI data acquisition and processing

MALDI MSI data were acquired at raster width of 80 µm using a 7T scimaX-MRMS mass spectrometer (Bruker Daltonics, USA) equipped with a SmartBeam II laser and a MALDI source. Peak-picked MALDI-IMS data were imported into SCiLS Lab 2024b software (SCiLS, Bremen, Germany) for image visualization and analysis. Compound identifications were assigned based on both accurate mass (<2 ppm mass accuracy) and isotope pattern matches to free source metabolite and lipid databases, including the Human Metabolome Database (HMDB), KEGG and LIPIDMAPs. Custom software written in Python programming language^71^ was used for mass spectrometry data processing. The software is available under GNU General Public License version 3 and is provided in Supplementary Data 2. Mean image intensities between corresponding brain regions were compared using two-tailed t-test.

### Statistical analysis

Data analysis was performed using R (version 3.6.0+) in RStudio (Version 24.12.1) or OriginPro (Version 9.8). All data are presented as mean ± s.d.. Intergroup differences between two groups were analyzed using unpaired two-tailed t-test or one-way ANOVA with Dunnett’s multiple comparisons test. For other group comparisons, multiple t-tests with false discovery rate (FDR) correction for multiple comparisons were applied. Statistically significant differences are indicatedas follows: *p<0.05, **p<0.01, ***p<0.001, ^§^p<0.0001. Number n indicates biological replicates (mitochondrial isolations or data obtained from individual mice).

## Acknowledgment

This work was supported by grants from the National Institutes of Health R01NS112381 and R01NS131322 (A. G.), R01NS100850, R01NS133383 (V.T.). Work on the Typhoon scanner was partially supported by NIH grant S10OD030335. I.W. was supported by the German Research Foundation (DFG #515944830) and SFB1531 (#456687919). We are grateful to Navdeep Chandel (Northwestern University, USA) for generously supplying us with the C57BL/6Rosa26^AOX-lsl/wt^ mouse strain used in this study. We also thank Howard Jacobs (Tampere University, Finland) for the kind gift of AOX antiserum. We are also grateful to Prof. Sergei Vinogradov (Department of Biochemistry and Biophysics, University of Pennsylvania) for fruitful discussion on *in vivo* oxygen measurements.

## Data availability

All data from the manuscript are available from the corresponding author on request. Proteomics data have been deposited in PRIDE under accession number PXD063555. Source data are provided with this paper.

## Code availability

All coding packages and custom software developed and used in this study are described in the Methods and are available from the corresponding author upon request.

## Author contributions

A.G. and V.T. conceived the project, designed research and analysed data. B.Y.S., S.S., F.A., I.G. Z.N. performed *in vitro* experiments. B.Y.S., P.Z and L.Z. carried out OGD-based assays. S.G. and Q.C. performed all metabolomics analyses. I.W performed BIAM proteomics. S.K., A.N. and C.K. advised on computational analysis. V.T., S.S., B.Y.S designed and performed *in vivo* mouse model of IR. B.Y.S., I.G., F.A., prepared samples and carried out enzymatic assays studies. C.K., A.N. and S.K. performed computational analysis. J.M., O.B., provided advice on mice breeding and contributed. A.A.S., G.M., S.S.G. provided advice on data analysis. G.M., M.S. and A.G. wrote the paper and integrated comments from the other authors.

## Extended Data

**Figure 1.**
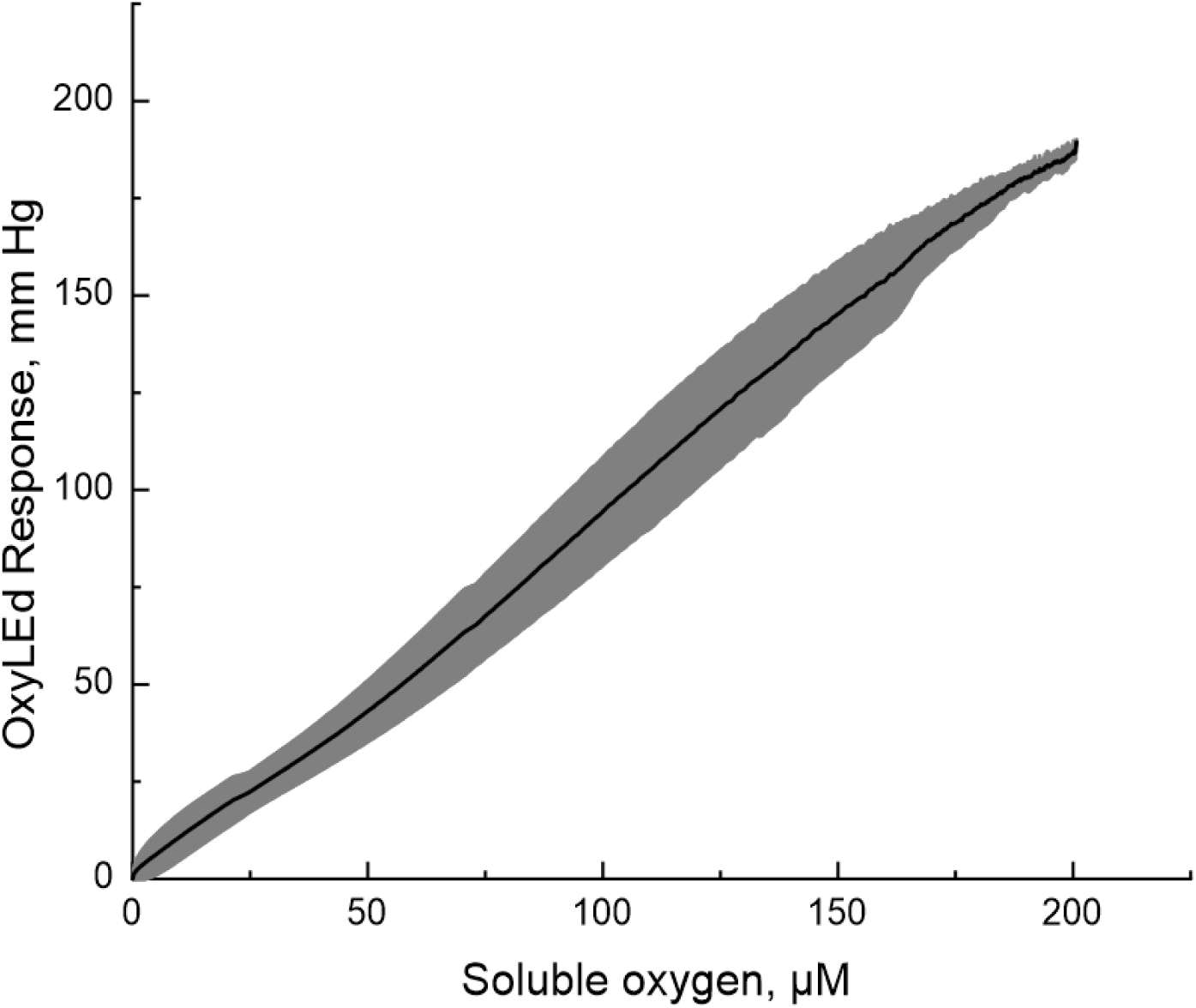
OxyPhore PG4 phosphorescence dye calibration for in situ oxygen measurement. OxyPhore. 3 µM of PG4 dye was added to 3 ml of PBS in the Oroboros respirometer chamber with phosphorescence decay time measuring Oxyled system fiberoptics attached to the glass wall window. To change oxygen levels, humidified pre-purified argon gas was equilibrated with the media of the respirometer chamber at a flow rate of 20 mL/min through 1 mL gas headspace above the liquid phase. This system allows simultaneous monitoring of oxygen concentration in the Oroboros chamber and measuring OxyLed signal change. The graph represents the average of three typical experiments where oxygen was replaced from the liquid phase and time decay of phosphorescence was measured *vs* soluble oxygen concentration (grey error bars represent s.d., n=3).

**Figure 2.**
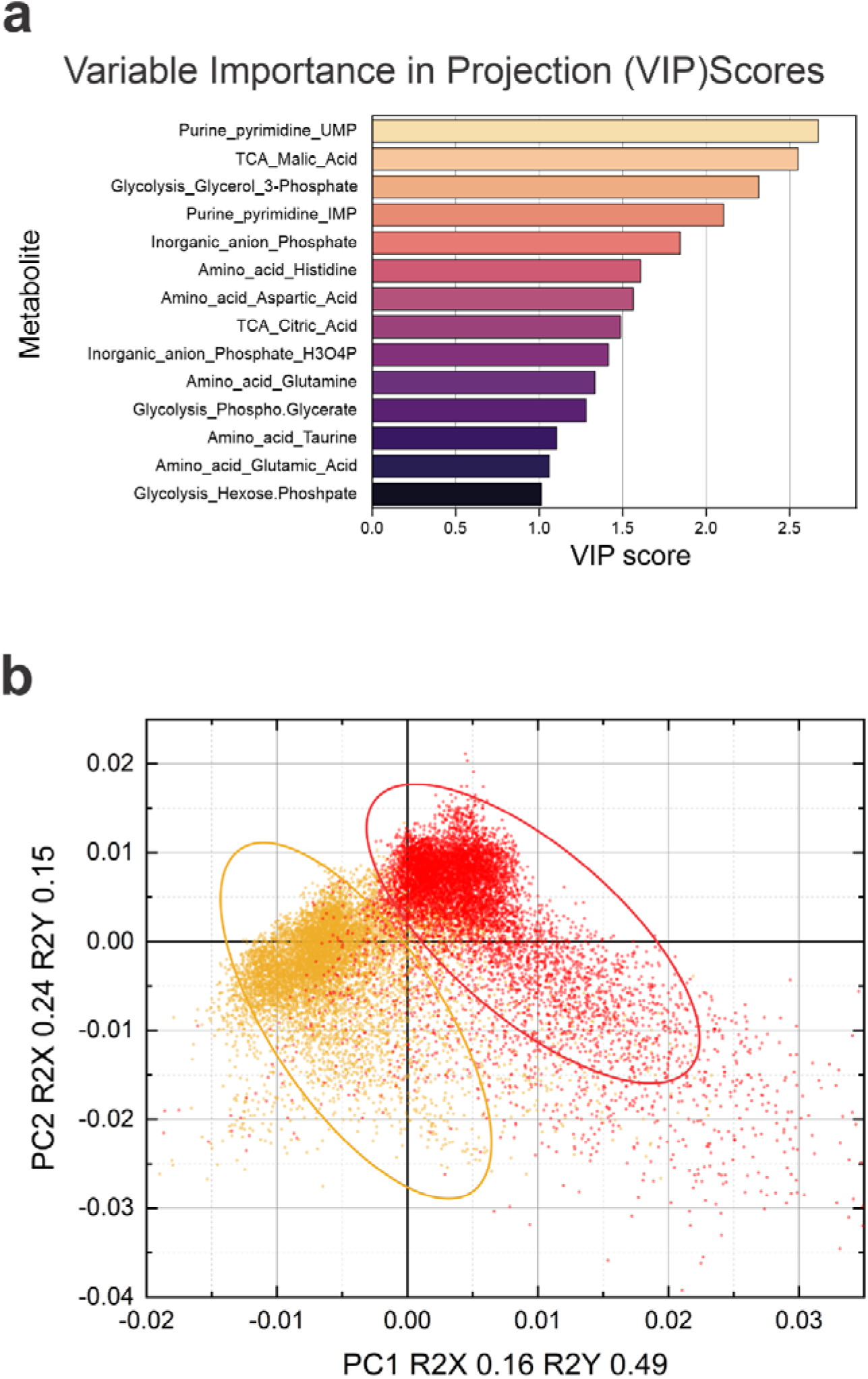
MALDI Imaging metabolomics analysis (relates to Fig. 3). **a**, variable Importance in Projection (VIP) scores from PLS-DA highlight the top metabolites that distinguish between healthy and ischemic regions in MALDI MSI data. **b**, PLS-DA scores illustrate clear separation of ischemic (red) and healthy (yellow) regions in the right and left hemispheres, respectively. Each pixel in the MALDI image was manually labelled as ischemic or healthy and was treated as an individual data point, with metabolites’ levels at this location used as features.

**Figure 3.**
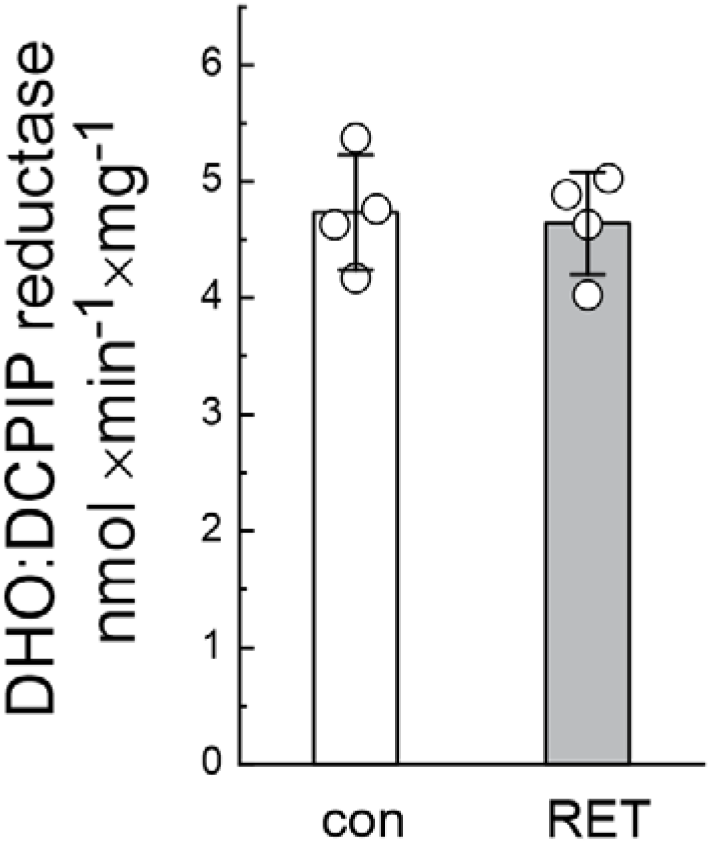
Dihydrorootate dehydrogenase (DHODH) activity is not affected by RET (relates to Fig 4). Activity was measured in intact mitochondria before (con) and after 20 min of oxidation of succinate (RET) as performed in Fig. 4. Aliquots were taken, and DHO:DCPIP reductase activity was assessed in permeabilized mitochondria as described in Materials and Methods section. Data are presented as mean ± s.d.; individual points represent biological replicates. Statistical significance was assessed by unpaired two-tailed t-test.

**Figure 4.**
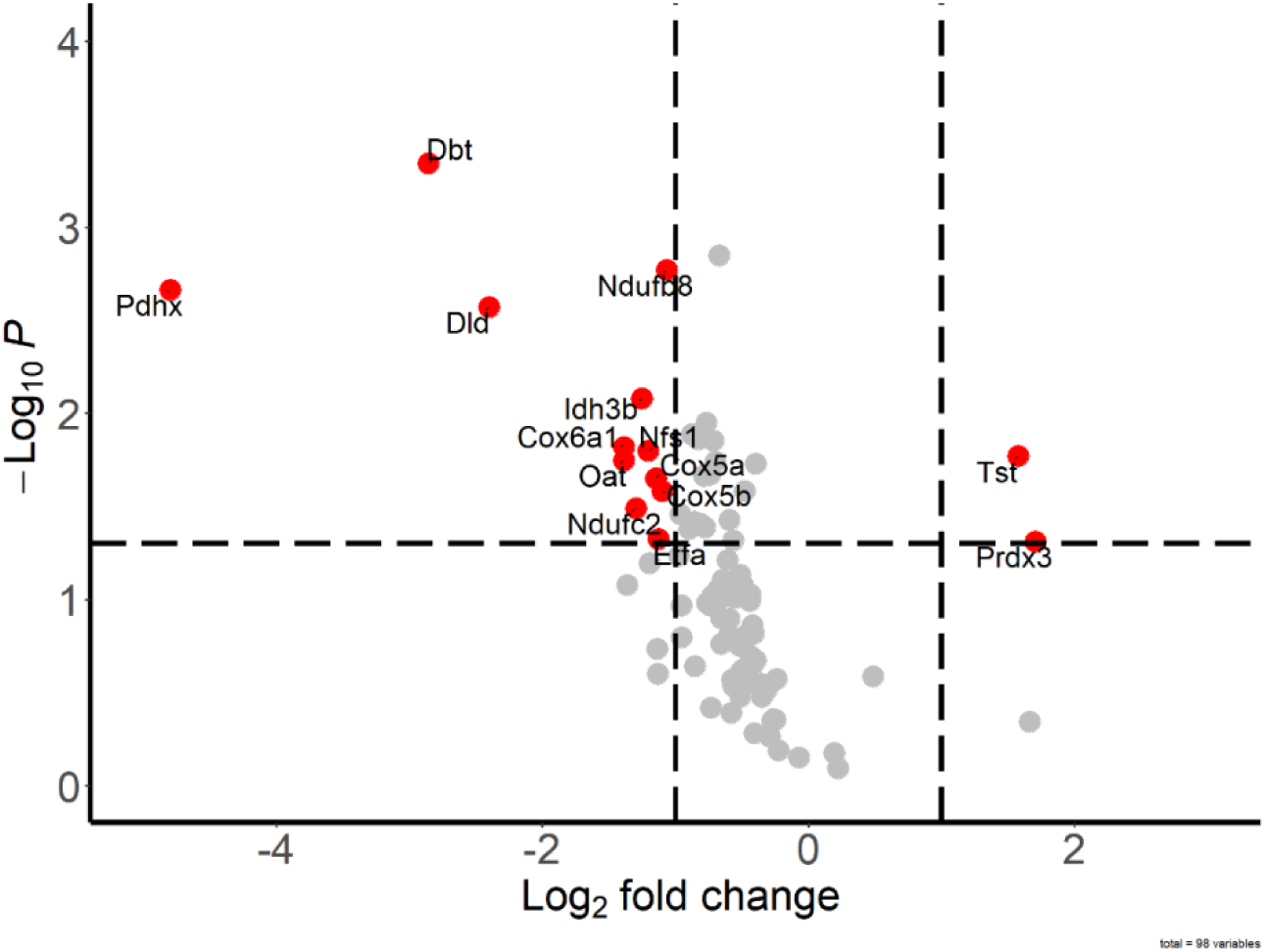
RET does not affect thiol redox state of complex I subunit around FMN-binding site. BIAM switch assay of intact brain mitochondria before and after RET for identification of oxidized proteins. Mitochondria from P10 mice were incubated with or without RET in the standard assay medium and 35 µg protein aliquots were taken before (control) and after 20 min incubation with 5 mM succinate (RET) (n=5 in each group). After protein precipitation MS-coupled BIAM switch assay was performed. Significant differentially oxidized proteins marked as red dots (p-value < 0.05; fold-change >2). The complete BIAM dataset can be found in Supplementary File 3.

**Figure 5.**
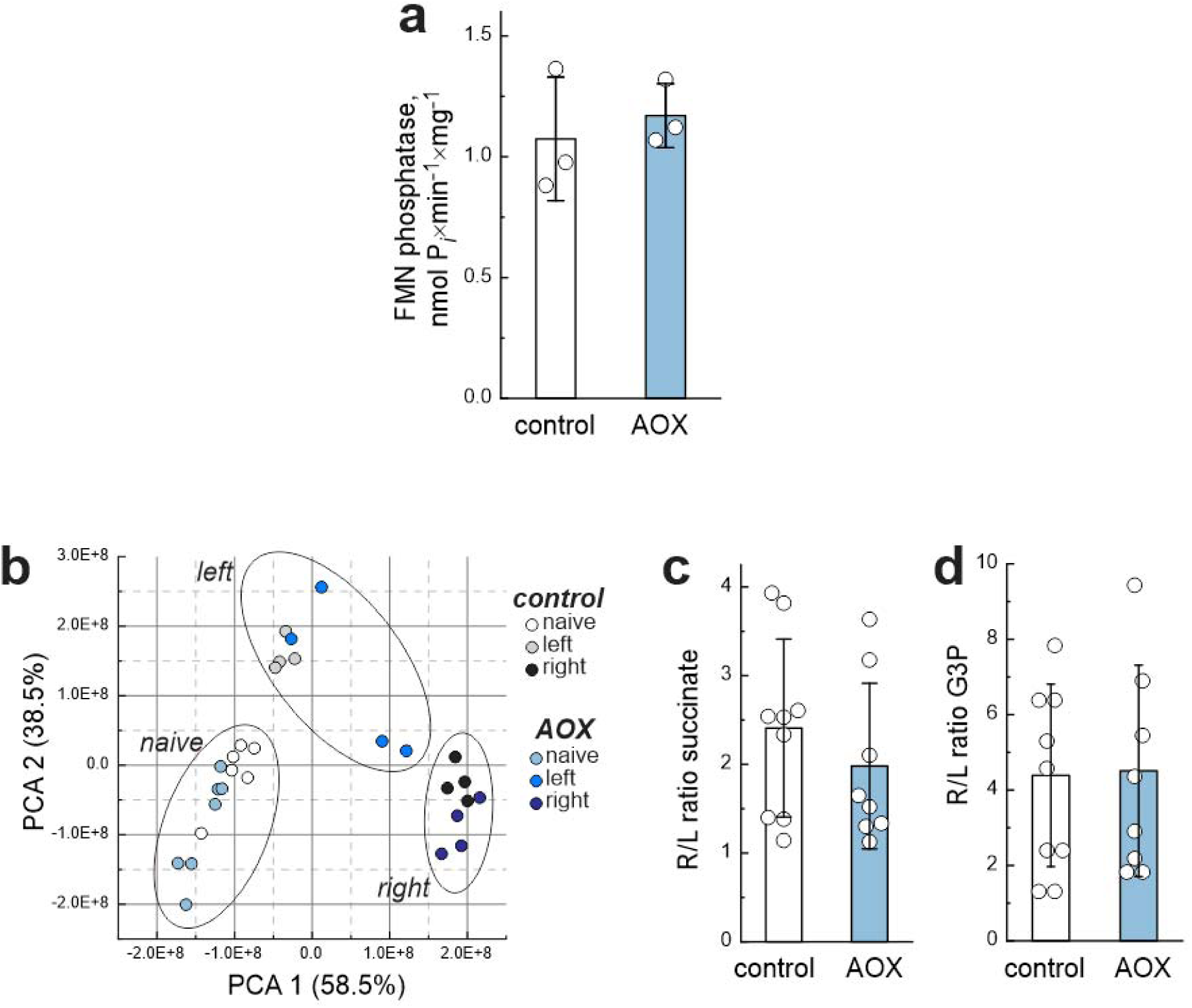
AOX expression does not affect mitochondrial metabolic profiles after HI/R (relates to Fig. 5). **a**, FMN phosphatase activity of matrix fraction isolated from intact brain mitochondria of control and AOX-expressing P10 mice. Inorganic phosphate release during FMN phosphatase reaction was measured as described in Ma erial and Methods**, b**, PCA of mitochondrial samples from control and AOX-expressing mice under naïve conditions and after *in vivo* HI measured in samples from left or right hemispheres. Each point represents an individual biological replicate. Ratio of RET metabolites; **c,** succinate and **d,** glycerol 3-phosphate levels between right and left hemispheres after *in vivo* HI. Data in **a, c, d** are presented as mean ± s.d.; individual points represent biological replicates. Statistical significance was assessed by unpaired two-tailed t-test. The complete metabolomics datasets can be found in Supplementary Table 4.

**Figure 6.**
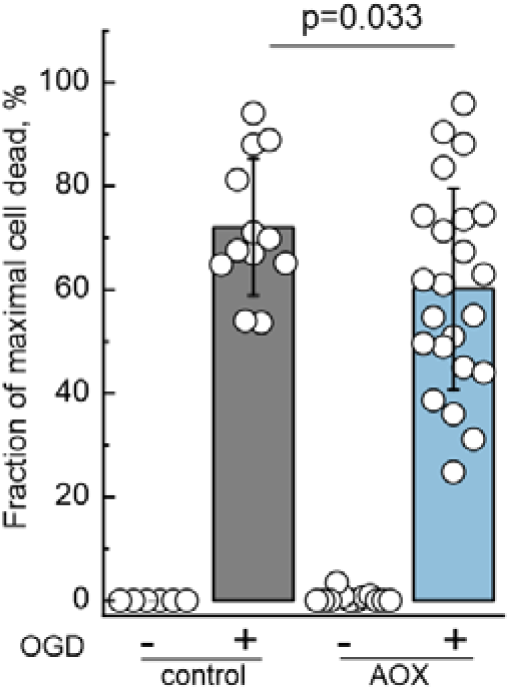
Quantitative analysis of cell death after OGD treatment (relates to Fig. 6). Data are presented as the percentage of propidium iodide (PI) fluorescence 24 h post-OGD. This analysis includes the subtraction of fluorescence prior to any treatment in control (grey bar), and AOX mice (blue bar). No PI-positive fluorescence was detected at baseline as indicated in the left bars for each genotype. Data are presented as mean ± s.d.; individual points represent biological replicates. Statistical significance was assessed by unpaired two-tailed t-test. n = 12 slices for control, and n = 24 slices for AOX.

**Figure 7.**
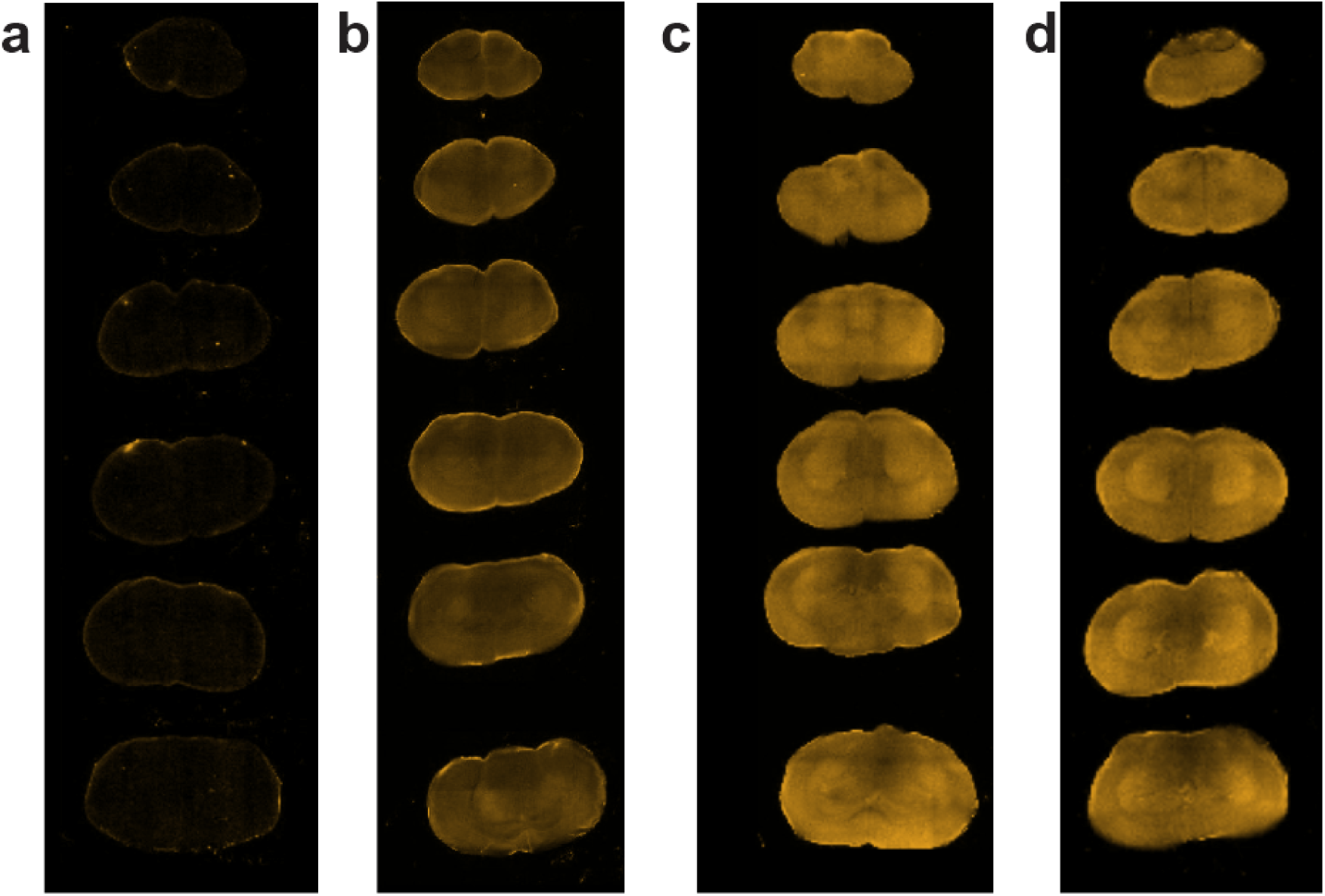
DHE oxidation as a measure of ROS production in brain slices from control and AOX mice. **a**, Basal tissue autofluorescence in SHAM brain mouse without DHE injection. Fluorescence measurements 4 h after 5 mg/kg of DHE intraperitoneal injection in **b,** naïve control animals or after IR injury in **c,** control, and **d,** AOX-expressing mice.

