## Supplemental Table 5 for "Rescuing Ischemic Brain Injury by Rewiring Mitochondrial Electron Flow" (from another version of this preprint: version 1, 2025-05-23)

List of reagents

| **Reagent** | **Manufacturer** | **Catalog number** |
| --- | --- | --- |
| Acetic acid (OmniTrace Ultra^TM^) | Millipore-Sigma | AX0078-7 |
| Acetonitrile (Optima, LC/MS) | Fisher Scientific | A9554 |
| Adenosine 5’-diphosphate monopotassium salt dihydrate, ADP | Sigma-Aldrich | A5285 |
| Alamethicin | Cayman Chemicals | 11425 |
| 9-aminoacridine, 9AA | Sigma-Aldrich | 90-45-9 |
| Ammonium acetate | Sigma-Aldrich | 372331 |
| Ammonium formate (LC/MS) | Sigma-Aldrich | 70221 |
| AmplexTM UltraRed Reagent | Thermo Fisher Scientific | A36006 |
| Antimycin A | Sigma-Aldrich | A8674 |
| Atpenin A5 | Cayman Chemicals | 11898 |
| 6-aminocaproic acid | Sigma-Aldrich | A2504 |
| N-Methyl-D-aspartic acid | Sigma-Aldrich | M3262 |
| BCA protein assay kit | Thermo Fisher Scientific | 23225 |
| Bovine serum albumin, fatty acid free | Sigma-Aldrich | A6003 |
| Brequinar | Cayman Chemicals | 24445 |
| Cytochrome C from equine heart | Sigma-Aldrich | C7752 |
| Carbonyl cyanide 4-(trifluoromethoxy)phenylhydrazone, FCCP | Sigma-Aldrich | C2920 |
| 2,6-Dichloroindophenol, DCIP | Sigma-Aldrich | D-1878 |
| Decylubiquinone, DBQ | Sigma-Aldrich | D7911 |
| Digitonin | Sigma-Aldrich | D141 |
| Dihydroethidium, DHE | Cayman Chemicals | 12013 |
| Dimethyl sulfoxide | Sigma-Aldrich | D8418 |
| n-Dodecyl β-D-maltoside, DDM | Sigma-Aldrich | D4641 |
| n-Dodecyl-β-D-maltoside, DDM | Cayman Chemicals | 16494 |
| DL-Dithiotreitol, DTT | Sigma-Aldrich | D0632 |
| L-Dihydroorotic acid | Sigma-Aldrich | D7128 |
| n-Ethylmaleimide, NEM | Sigma-Aldrich | 04259 |
| Essentially fatty acid–free bovine serum albumin, BSA | Sigma-Aldrich | A6003 |
| Ethylene glycol-bis(2-aminoethylether)-N,N,N′,N′-tetraacetic acid, EGTA | Sigma-Aldrich | E3889 |
| Ethylenediaminetetraacetic acid tetrasodium salt hydrate, EDTA | Sigma-Aldrich | ED4S |
| L-Glutamic acid | Sigma-Aldrich | 49449 |
| Glycerol | Sigma-Aldrich | G5516 |
| rac-Glycerol-1-phosphate sodium salt, G3P | Sigma-Aldrich | 61668 |
| Hexaammineruthenium (III) chloride | Sigma-Aldrich | 262005 |
| 4-(2-Hydroxyethyl)piperazine-1-ethanesulfonic acid, N-(2-Hydroxyethyl)piperazine-N′-(2-ethanesulfonic acid), HEPES | Sigma-Aldrich | H4034 |
| Hydrogen peroxide solution, H_2_O_2_ | Sigma-Aldrich | 216763 |
| HRP-antirabbit, Peroxidase AffiniPure® Goat Anti-Rabbit IgG (H+L) | Jackson ImmunoResearch | 111-035-003 |
| Imidazole | Sigma-Aldrich | 792527 |
| Indium Tin Oxide, ITO | Delta Technologies | CB-90IN-S111 |
| Magnesium chloride solution | Sigma-Aldrich | M1028 |
| Malachite green phosphate assay kittHERMO f | Sigma-Aldrich | MAK307 |
| L-Malic acid disodium salt monohydrate | Sigma-Aldrich | 233935 |
| Methanol (Optima^TM^ LC/MS) | Fisher Scientific | A4564 |
| N-(1-naphthyl)ethylenediamine dihydrochloride, NEDC | Sigma-Aldrich | 1465-25-4 |
| SF-6847 | Sigma-Aldrich | T182 |
| Sodium malonate dibasic monohydrate | Sigma-Aldrich | M4795 |
| Mannitol | Sigma-Aldrich | 63559 |
| NativePAGE^TM^ Bis-Tris Mini Protein Gels, 3 to 12% | Thermo Fisher Scientific | BN1001BOX |
| β-Nicotinamide adenine dinucleotide, reduced disodium salt hydrate, NADH | Sigma-Aldrich | N8129 |
| Nitro Blue Tetrazolium (chloride) | Cayman Chemicals | 17341 |
| PBS | Thermo Fisher Scientific | 10010049 |
| Ponceau S | Thermo Fisher Scientific | 161470100 |
| Peroxidase from horseradish, HRP | Sigma-Aldrich | P8250 |
| Potassium chloride | Sigma-Aldrich | P3911 |
| Potassium cyanide | Sigma-Aldrich | 207810 |
| 2-Propanol | Fisher Scientific | A461-500 |
| Propidium iodide | Thermo Fisher | P3566 |
| Q1, coenzyme Q1 | Sigma-Aldrich | C7956 |
| Sodium pyruvate | Sigma-Aldrich | P5280 |
| Riboflavin 5’-monophosphate sodium salt hydrate, FMN | Sigma-Aldrich | F2253 |
| Rotenone | Sigma-Aldrich | R8875 |
| Sodium deoxycholate | Sigma-Aldrich | D6750 |
| Sodium dodecyl sulfate, SDS | Sigma-Aldrich | L3771 |
| Sodium succinate dibasic | Sigma-Aldrich | 14160 |
| Sucrose | Sigma-Aldrich | 84097 |
| Superoxide Dismutase bovine, SOD | Sigma-Aldrich | S9697 |
| 2, 3, 5-Triphenyltetrazolium chloride, TTC | Sigma-Aldrich | T8877 |
| Tricine | Sigma-Aldrich | T0377 |
| Trichloroacetic acid | Sigma-Aldrich | 91228 |
| Trizma® base, Tris | Sigma-Aldrich | T1503 |
